# From Chemoproteomic-Detected Amino Acids to Genomic Coordinates: Insights into Precise Multi-omic Data Integration

**DOI:** 10.1101/2020.07.03.186007

**Authors:** Maria F. Palafox, Valerie A. Arboleda, Keriann M. Backus

## Abstract

The integration of proteomic, transcriptomic, and genetic-variant annotation data will improve our understanding genotype-phenotype associations. Due, in part, to challenges associated with accurate inter-database mapping, such multi-omic studies have not extended to chemoproteomics, a method that measure the intrinsic reactivity and potential ‘druggability’ of nucleophilic amino acid side chains. Here, we evaluated two mapping approaches to match chemoproteomic-detected cysteine and lysine residues with their genetic coordinates. Our analysis reveals that databases update cycles and reliance on stable identifiers can lead to pervasive misidentification of labeled residues. Enabled by this examination of mapping strategies, we then integrated our chemoproteomic data with *in silico* generated predictions of genetic variant pathogenicity, which revealed that codons of highly reactive cysteines are enriched for genetic variants that are predicted to be more deleterious. Our study provides a roadmap for more precise inter-database comparisons and points to untapped opportunities to improve the predictive power of pathogenicity scores and to advance prioritization of putative druggable sites through integration of predictions of pathogenicity with chemoproteomic datasets.

## INTRODUCTION

Understanding how proteins work is the bedrock of functional biology and drug development. The identification of amino acids that directly regulate a protein’s activity (e.g. catalytic residues, residues that drive interactions, or residues important for folding or stability) is an essential step to functionally characterize a protein. Delineation of amino acid-specific functions is typically accomplished using site-directed mutagenesis (Starita et al. 2015; Hemsley et al. 1989). While such studies can identify functional hotspots in human proteins, they are typically limited in scope and largely restricted to proteins easily expressed *in vitro*. With the advent of next generation sequencing and CRISPR-based mutagenesis, deep mutational analysis can now be scaled to individual genes (e.g. *TP53* and *BRCA1*) (Starita et al. 2015; Boettcher et al. 2019), but such studies have not been extended genome-wide.

This problem of identifying the function of a specific amino acid parallels one of the central challenges of modern genetics: interpretation of the pathogenicity of the millions of genetic variants found in an individual’s genome. Many *in silico* algorithms, such as M-CAP (Jagadeesh et al. 2016), CADD (Kircher et al. 2014), PolyPhen (Adzhubei et al. 2010), and SIFT (Vaser et al. 2016) integrate data such as sequence conservation, metrics of sequence constraint, and other functional annotations to provide a metric for stratification of benign and pathogenic variants. In the absence of experimental data, these scores provide a quantitative metric to rank genetic variants for their effect on a phenotype, something particularly important in the era of genome-wide association studies and sequencing studies.

Beyond genetic variation, a frequently overlooked parameter that defines functional ‘hot-spots’ in the proteome is amino acid side-chain reactivity, which can fluctuate depending on the residue’s local and 3-dimensional protein microenvironment. Mass spectrometry-based chemoproteomics methods have been developed that can assay the intrinsic reactivity of thousands of amino acid side chains in native biological systems (Weerapana et al. 2010; Backus et al. 2016; Hacker et al. 2017). Using these methods, previous studies, including our own, revealed that “hyper-reactive” or pKa perturbed cysteine and lysine residues are enriched in functional pockets. These chemoproteomic methods can even be extended to measure the targetability or “druggability” of amino acid side chains, which has revealed that a surprising number of cysteine and lysine side chains can also be irreversibly labeled by small drug-like molecules (Weerapana et al. 2010; Backus et al. 2016; Hacker et al. 2017). Complicating matters, for the vast majority of these chemoproteomic-detected amino acids **(CpDAA)**, the functional impact of a missense mutation or chemical labeling remains unknown. Integrating chemoproteomic data with genomic-based annotations represents an attractive approach to stratify amino acid functionality and to identify therapeutically relevant disease-associated pockets in human proteins.

Such multi-omic studies require mapping a protein’s sequence back to genomic coordinates, through the transcript isoforms, in essence reverse engineering the central dogma of molecular biology. Accurate mapping between genomic coordinates and amino acid positions remains particularly challenging, due in part to the diversity of cell-type specific transcript and protein isoforms and the non-linear relationship between genomic, transcriptomic, and protein databases. One approach to address these challenges is through proteogenomics, where custom FASTA files are generated from whole exome or RNA-sequencing data. However, such approaches are not typically scalable or cost-effective. Furthermore, many proteomic datasets, particularly previously acquired and public datasets, lack matched genomic data, precluding proteogenomic analysis.

Many computational tools have been developed for inter-database mapping, including using unique identifiers (Agrawal and Prabakaran 2020; L. M. Smith et al. 2019; Durinck et al. 2009), methods to map genomic coordinates to protein sequences and structures(Stephenson et al. 2019; Sehnal et al. 2017; Sivley et al. 2018; David and Yip 2008), and tools for codon-centric-based annotation of genetic variants (Gong et al. 2014; Schwartz et al. 2019). One key application of these tools is in the improved prediction of variant pathogenicity (Guo et al. 2017). Complicating such data integration, while many predictive genetic scores are built on the GRCh37 reference genome (frozen in 2014), the proteomic UniProtKB reference is based on GRCh38. Further complicating matters, the unsynchronized and frequent updates to widely used databases, such as UniProtKB and Ensembl, results in a constantly evolving landscape of genome-, transcriptome- and proteome-level sequences and annotations, which further confounds multi-omic data integration, particularly for residue level analyses.

Focusing on previously identified CpDAAs (Weerapana et al. 2010; Backus et al. 2016; Hacker et al. 2017), we first quantitatively assess the impact of mapping methods, including the use of stable and versioned identifiers and tools for residue-coordinate mapping, on the fidelity of data integration. We then apply an optimized mapping strategy to annotate CpDAA positions with predictions of genetic variant pathogenicity. Our study uncovers key sources of inaccurate mapping and provides fundamental guidelines for multi-omic data integration. We also reveal that highly reactive cysteines are enriched for genetic variants that have high predicted pathogenicity (high deleteriousness), which supports both the utility of predictive scores to further power proteomics datasets and the use of chemoproteomics to add another layer of interpretation to missense genetic variants. As many databases move to GRCh38, we anticipate that our findings will provide a roadmap for more precise inter-database comparisons, which will have wide ranging applications for both the proteomics and genetics communities.

## RESULTS

### (1) Characterizing the dynamic mapping landscape relevant to CpDAA data integration

Our first step to achieve high fidelity multi-omic data integration was to establish a comprehensive set of test data. For this we turned to publicly available chemoproteomics datasets (Backus et al. 2016; Hacker et al. 2017; Weerapana et al. 2010). To generate our test dataset, we first aggregated the CpDAAs identified in all three prior studies and then confirmed that all UniProtKB IDs and residue numbers were valid (**Table S1**). All three original CpDAA datasets had been previously searched against the same reference proteome (November 2012 UniProtKB release), a non-redundant FASTA file of protein sequences identified by UniprotKB stable IDs. In these datasets, each detected amino acid is represented as the associated stable identifier (**Table S2**) and amino acid position based on the canonical sequence (e.g. P04637_C141 represents cysteine 141 in UniProtKB ID P04637 sequence, which is TP53). After data merge and ID validation (see Methods), we recovered 99% of the original chemoproteomic datapoints, with a small fraction lost due to absence of protein stable identifiers in the 2012 UniProtKB FASTA file or mis-matched coordinates for detected cysteine and lysine. In aggregate, our dataset consists of 6,404 identified cysteine residues and 9,213 identified lysine residues within 4,084 unique proteins. These 15,617 CpDAAs are further sub-categorized by the measures of amino acid intrinsic reactivity and potential targetability.

As our overarching objective was to characterize CpDAAs using functional annotations based on the protein, transcript, or gene coordinates (**Figure 1A**), our next step was to map the CpDAA UniprotKB stable identifiers to the identifiers of corresponding proteins, genes, and transcripts in more recent versions of UniProtKB, Ensembl and GENCODE. We chose to map our data using stable identifiers, instead of versioned identifiers (**Table S2**), for two reasons. First, the CpDAAs were identified by stable, but not versioned, identifiers. Second, stable identifiers offer the seeming advantage of remaining constant through update cycles. However, one particularly confusing and notable aspect of the stable identifier is that the word “stable” in this context does not mean permanent, reliable, or immutable. Specifically, the associated sequence linked to a stable identifier can change over database version releases. These changes complicate residue-level mapping across different database releases and can cause residue mismapping where the amino acid position is correct in one release but incorrect in a future updated version of the database. Therefore, we evaluated established methods for inter-database mapping, including <u>ID mapping</u>, <u>residue-residue mapping</u>, and <u>residue-codon mapping</u> (See **Table S2** for detailed descriptions of each type of mapping).

**Figure 1.**
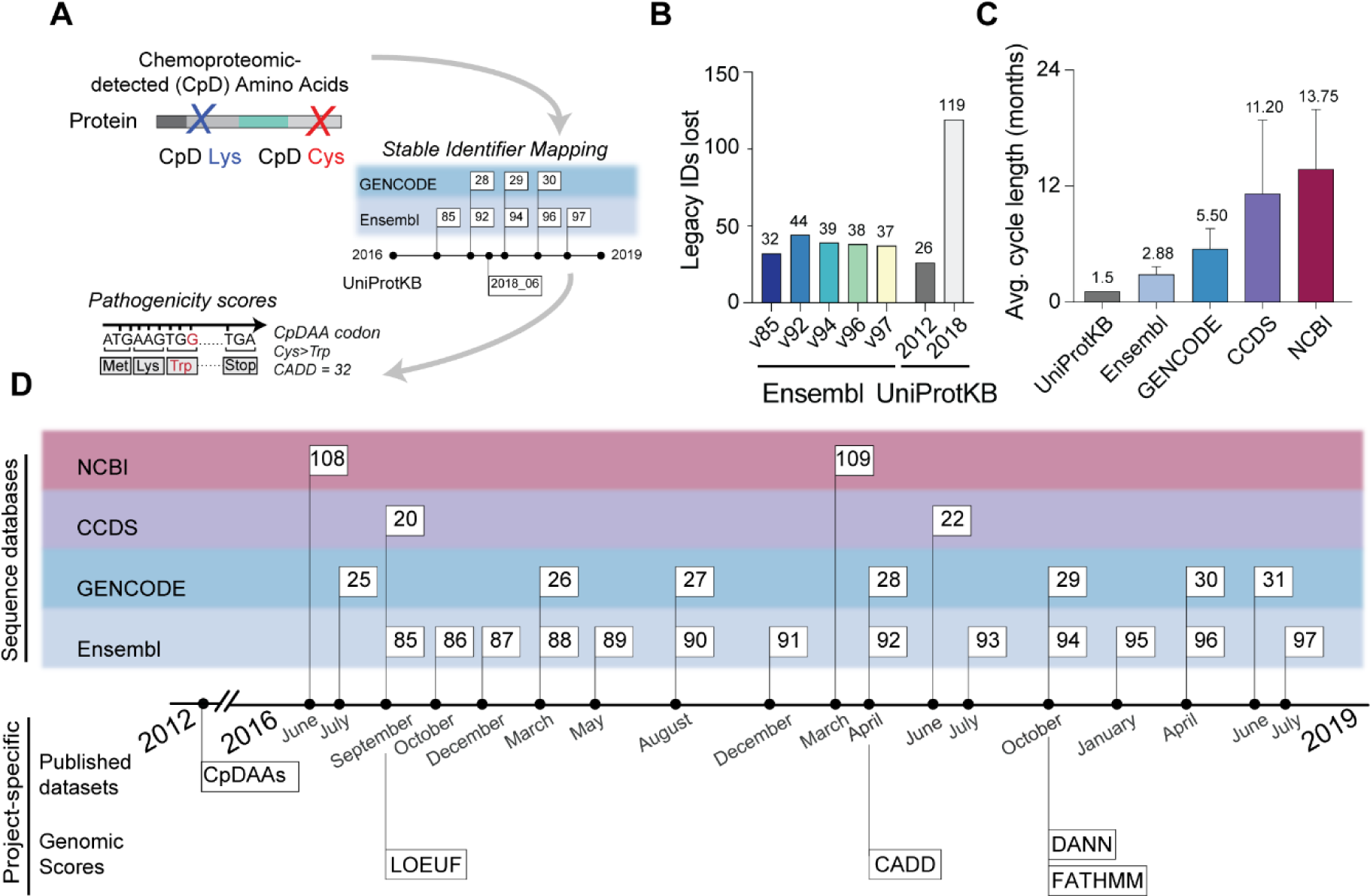
Stable identifier mapping across database releases. A) Overall objective of this study is to use stable identifiers to map chemoproteomic detected amino acids (CpDAAs), denoted here as red/blue ‘X,’ to variant pathogenicity scores. B) Shows the number of stable UniProtKB protein IDs from cysteine and lysine chemoproteomics studies in original legacy chemoproteomics dataset (4,119 Uniprot stable IDs in aggregate)(Hacker et al. 2017; Backus et al. 2016; Weerapana et al. 2010) that fail to map to IDs in more recent releases of Ensembl and UniProtKB. C) Average update cycle length across major gene annotation databases. D) Timeline of gene annotation database releases, including Ensembl releases tested for compatibility (**Figure 3**) to CpDAA coordinates based on canonical UniProtKB protein sequences.

We collected a test set of Ensembl releases (**Table S3**) to benchmark the fidelity of specific mapping methods, using our CpDAA dataset as a starting point. Specific releases were prioritized to highlight how specific updates and integrations impacted data mapping. Specific releases were prioritized that (1) represented reference releases based on the GRCh37 or GRCh38 reference genome, (2) were compatible with the latest Consensus CoDing Sequence (CCDS) update for the human genome (release 22), (3) were used in dbNSFPv4.0a and CADDv1.4, two resources that integrate functional annotations for all possible non-synonymous single nucleotide variants (SNV) (Kircher et al. 2014; Liu et al. 2016), and (4) were associated with a commonly used version of the Ensembl Variant Effect Predictor (VEP) (McLaren et al. 2016).

With this data in hand, we next tracked the loss of proteins and CpDAAs during *intra-*database mapping of UniProtKB releases and *inter-*database mapping of multiple Ensembl releases to a frozen release of the UniProtKB human proteome. Through this analysis we sought to quantify how database update-induced changes to the protein sequences associated with stable identifiers would affect CpDAA mapping. Gratifyingly, only a handful of the original 4,119 IDs were lost due to database updates, both for Ensembl and UniProtKB (**Figure 1B**). While all Ensembl datasets showed similar losses, Ensembl v85 modestly outperformed more recent versions, consistent with the v85 release date being closest in time to the UniProtKB release on which legacy data was based. The greatest identifier loss was observed from mapping UniProtKB-based legacy data to the UniProtKB-SwissProt curation of the human proteome from 2018, with 119 entries not found in the 2018 dataset. We ascribe this identifier loss to both UniProtKB updates and to the higher-level of curation for proteins in the 2018 dataset, which includes only Swiss-Prot canonical protein sequences with an entry term in the CCDS database cross-reference (xref) files. Of note, CCDS gene IDs are manually reviewed and linked to UniProtKB-SwissProt. In contrast, other database cross-references include protein IDs that are not manually reviewed. Both the manually curated and automatically generated protein IDs are included in the TrEMBL database, which, as a result is comprised of a significantly larger sets of UniProtKB IDs, when compared to CCDS (**Figure EV1**).

Accurate residue-level mapping between sequences from different database sources is further complicated by the frequent and unsynchronized update cycles of independent databases. Therefore, we next sought to identify the optimal releases for data mapping. We assembled a comprehensive inventory of all database updates for Ensembl, GENCODE, NCBI, and CCDS (**Table S4**) between the dates of August 2013 - July 2019 (6 years). In total, we noted the release dates of 25 Ensembl, 13 GENCODE, 6 CCDS (*homo sapiens* only), and 5 NCBI releases. Quantification of the average update cycle for each database across this time period revealed that UniProtKB has the shortest mean update cycle length (∼6 weeks) (**Figure 1C**). In contrast, NCBI is only updated yearly. Many updates are not synchronized (**Figure 1D**), which can create a lag between versions of databases used to create identifier cross-reference (xref) files. For example, stable ID mapping files provided by Ensembl for UniProtKB proteins may not share identical sequences if not used within the short 4-week window between UniProtKB updates. These changes can accumulate over time and can lead to data loss and residue mis-annotation. From this analysis, we concluded that using the CCDS UniProtKB release was optimal for integration with multiple predictions of pathogenicity, including CADD (Kircher et al. 2014), DANN (Quang, Chen, and Xie 2015), and FATHMM (Rogers et al. 2018).

### (2) Updates to canonical sequences assigned to UniProtKB stable identifiers can lead to intra-database mismapping of CpDAAs

Proteomics datasets, including the published CpDAA datasets, are routinely searched against FASTA files containing all canonical sequences (**Table S2**) of proteins encoded by the genome, identified by UniProtKB stable IDs. The use of canonical sequences, instead of all isoforms, substantially reduces the redundancy and complexity of database searches. An additional advantage of these stable IDs is that they remain unchanged across releases, which can facilitate rapid intra-database cross referencing. Therefore, our next step was to use these stable IDs to map between two UniProtKB releases (2012 and 2018). By mapping our data to the 2018 UniProtKB CCDS release, we aimed to take advantage of the extensive array of tools that have been developed to facilitate forward and reverse annotation between gene, transcripts, and protein coordinates (**Table S5**). Updating to the 2018 release was a requisite step for using these tools, as they overwhelmingly require recent cross-reference files. We performed residue-residue mapping—defined as a one-to-one correspondence between amino acids in proteins from different databases or release dates (**Table S2**)—of all CpDAA positions between the two versions of UniProtKB. We matched the canonical sequences that were linked with the 2012 stable ID vs the 2018 stable ID from our CpDAA data (**Table S6**). Our analysis showed the loss of 121 protein IDs, with 108 simply not found and the remaining 13 found to have different canonical sequences, resulting in mismapping or loss of the originally identified CpDAA residues.

The high concordance between these two UniProtKB releases, separated by six years, indicates that for the vast majority of UniProtKB updates, differences in release date should not complicate annotating legacy proteomics data with metrics based on more recently released gene, transcript, and protein sequences. However, we were surprised to find that several widely studied proteins, including protein arginine N-methyltransferase 1 (PRMT1, Q99873) (Tang et al. 2000), serine/threonine-protein kinase, (SIK3; Q9Y2K2) (Walkinshaw et al. 2013), and Tropomyosin alpha-3 chain (TPM3, P06753) were not found during intra-UniProtKB mapping (**Table S6**). We observed two main reasons for these losses: 1) changes to the canonical sequence associated with the UniProtKB stable ID and 2) changes to which isoform is assigned as the canonical sequence. While both the 2012 and 2018 sequences of PRMT1 are associated with UniProtKB stable ID Q99873, the 2018 sequence contains an additional short N-terminal sequence, not present in the 2012 sequence (**Figure 2A**). As a result, all 13 PRMT1 CpDAAs failed to map to the 2018 UniProtKB release. In the 2012 release of UniProtKB, the canonical sequence of the Peptidyl-prolyl cis-trans isomerase FKBP7 is associated with the versioned (isoform) ID Q9Y680-1, whereas in the 2018 release the canonical sequence is associated the versioned (isoform) ID Q9Y680-2, which lacks a short sequence (AAΔ125:162) in the middle of the protein. For FKBP7, this update fortuitously does not result in loss of CpD Lys83, as it is located n-terminal to the deletion. However, as showcased by PRMT1, updates to a protein’s canonical sequence, such as these, can easily result in data loss and mismapping.

**Figure 2.**
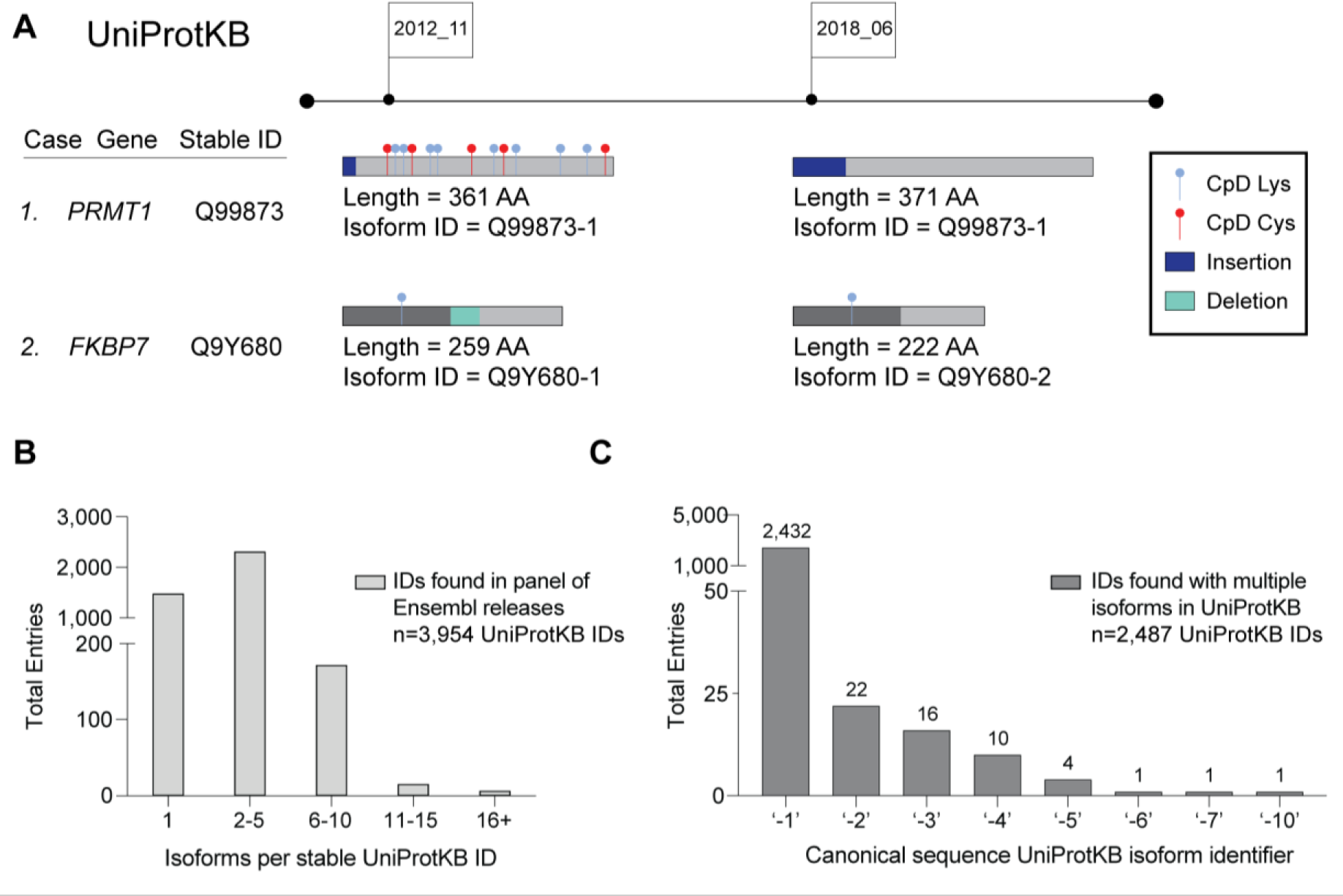
Stable and versioned identifier mapping between UniProtKB releases. A) Residue mismapping due to UniProtKB database updates to sequences associated with stable identifiers. Shown are two representative mapping outcomes. For PRMT1, addition of 10 amino acids at the protein N-terminus results in incorrect mapping of all 13 CpDAAs. For FKBP7, CpDAA Lys83 is correctly mapped in the 2018 release, as the sequence update occurs C-terminal to the modified residue. B) The number of isoforms associated with CpDAA UniProtKB stable IDs. C) Canonical isoform ID number associated with UniProtKB entries. Only CpDAA stable IDs with >1 isoform sequence were used for analysis.

These updates are, in essence, masked by the stable IDs, which do not flag sequence updates or changes to which isoform sequence is assigned as the canonical. Exemplifying this problem, we identified 45 stable identifiers with non-identical canonical protein sequences in the 2012 and 2018 UniProtKB releases (**Table S7**). Therefore, our next step was to investigate how the presence or absence of protein isoforms would affect intra-database mapping. Analysis of all protein entries associated with our CpDAA dataset revealed 58% of protein stable IDs have between 2-5 associated isoform sequences (**Figure 2B**). Some proteins had sequence variability with five proteins having more than 16 isoforms and Catenin delta-1 protein (CTNND1, O60716) had 32 isoforms, which was the greatest number of isoforms in our dataset (**Table S8**). Protein entries with only one isoform accounted for 37% of protein stable IDs (n=1,466).

Protein isoforms are identified by the ‘-X’ after the UniProtKB ID, where X represents the isoform number. A standard assumption of most mapping tools and proteomics databases is that the ‘-1’ sequence is the canonical sequence. However, a key finding from our isoform analysis is that the canonical sequence does not actually always correspond to the ‘-1’ isoform ID provided by UniProtKB. For 55 proteins (∼2%) of our CpDAA-containing proteins, the canonical sequence is not the ‘-1’ isoform (**Figure 2C** and **Table S9**). Strikingly, the canonical sequence can even be the ‘-10’ isoform, as is the case for the Ras-associated and pleckstrin homology domains-containing protein (RAPH1, Q70E73). In the context of database mapping, all of these non-‘-1’ canonical entries will likely result in mismapping using established tools.

### (3) Accurate residue-level mapping between UniProtKB and Ensembl is dependent on database update cycles

To investigate *inter-database mapping*, we next turned to ID cross-reference files **(Table S2)** that are released by Ensembl and UniProtKB. Cross-reference files can be used to convert between UniProtKB and Ensembl ID types. In the ideal scenario for inter-database mapping, we would have a 1:1 relationship between UniProtKB protein ID and amino acid position and Ensembl protein, transcript, and gene IDs and corresponding genomic coordinates. However, in practice, three major challenges arise with ID cross-referencing: 1) when cross reference stable IDs match, but corresponding sequences are not identical, 2) *multi-mapping*, where a UniProtKB ID maps to many Ensembl protein, transcript, and gene IDs, and 3) when the origin, both the time of releases and which database provided cross reference files, determines the mapping accuracy of datasets.

Sequence updates associated with a stable ID can lead to mismapping of gene-, transcript-, and protein-level annotations for CpDAAs. Glucose-6-phosphate dehydrogenase (G6PD, P11413) exemplifies how Ensembl IDs change with database updates from v85 to v92 (**Figure 3A**). For G6PD, the same UniProtKB ID maps to multiple Ensembl stable protein IDs (colored squares). Additionally, the protein sequence associated with specific Ensembl stable IDs varied across Ensembl releases. In fact, over time the same protein sequence was associated with four different Ensembl protein IDs with identical sequences (see first row in ‘Identical’) as well as three different Ensembl protein IDs with non-identical sequences (see second row in ‘Non-identical’). For G6PD, this significant redundancy at the protein ID level also extends to the Ensembl stable transcript and gene IDs (**Figure 3B**).

**Figure 3.**
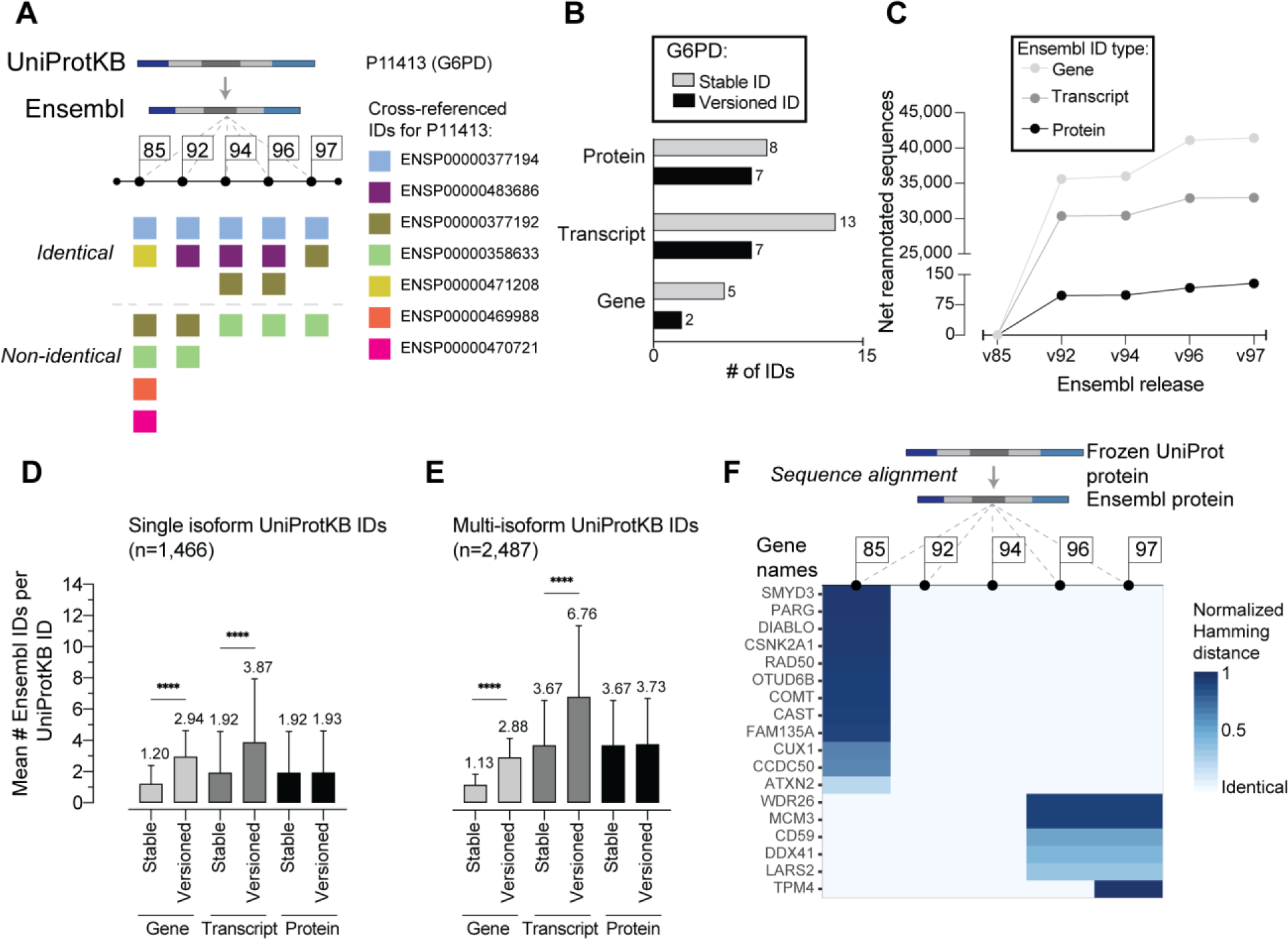
Mapping between UniProtKB stable IDs and Ensembl stable and versioned IDs across releases. A) CpDAA UniProtKB stable IDs map to multiple Ensembl stable protein IDs. The protein glucose-6-phosphate dehydrogenase (G6PD, UniProtKB ID P11413) maps to multiple Ensembl stable protein IDs including identical and non-identical sequences across all five Ensembl releases investigated. B) Number of stable and versioned Ensembl gene, transcript and protein IDs for G6PD across all five Ensembl releases shown in ‘A.’ C) Cumulative sequence re-annotations for Ensembl gene, transcript, and protein IDs since the v85 release. D-E) Average number of Ensembl gene, transcript, and protein IDs for (D) single isoform (n=1,466) and (E) multi-isoform (n=2,487) CpDAA UniProt entries. F) Heatmap depicts protein alignment scores by normalized Hamming distance, with 0 indicating no difference, comparing CpDAA UniProtKB protein sequences (n=3,953) to cross-referenced IDs of Ensembl protein sequences (n= 29,450) across the five Ensembl releases. For D-E, bar plots represent mean values ± SD for the number of Ensembl IDs per stable UniProt ID. Statistical significance was calculated using an unpaired Student’s T-test, **** p-value <0.0001.

Building upon the substantial number of Ensembl sequence updates observed for G6PD, we next assessed how pervasive sequence updates were for associated stable identifiers across all Ensembl gene, transcript, and protein IDs mapped to our set of CpDAA UniProtKB IDs. We counted the sequence version increments for each Ensembl stable ID, starting with the v85 release. The number of version updates varies for gene, transcript, and protein sequences associated with Ensembl IDs. We find that genes undergo the highest frequency of sequence re-annotation (**Figure 3C**). For example, between Ensembl v85 and v97, there are only 128 sequence updates for protein IDs compared with 32,949 sequence updates for transcript IDs and 41,439 for gene IDs (**Table S10)**. Therefore, this high variability in sequence updates for stable IDs make mapping using only stable IDs a challenging problem.

A solution to deciphering which of these many sequences are non-redundant, is to use cross reference files with Ensembl versioned IDs (**Table S2**), which adds an increment to a stable ID (i.e. ‘.1’ to ‘.2’) to indicate that a change to the associated sequence has occurred. For Ensembl stable identifiers, version number increments signify updates made to the associated amino acid or nucleotide sequence or to exon boundaries (“Ensembl Stable IDs” n.d.; Ruffier et al. 2017). These versioned IDs can reduce the many-to-one, one-to-many, or many-to-many relationships between gene, transcript, and protein IDs. For example, for protein tropomyosin alpha-4 chain (TPM4, P67936), during the update from v96 to v97, the stable protein identifier was showed version change from ‘.3’ to ‘.4’ (ENSP00000300933.3 to ENSP00000300933.4) which corresponds to a difference of 165 amino acids in the primary sequence (**Table S11a**). While versioned IDs appear to solve many mapping issues, they are challenging to work with as many of the cross-reference files made available by each database only provide stable IDs (and not versioned identifiers) for the database being cross-referenced.

Multi-mapping, where a single UniProtKB ID maps to many Ensembl protein, transcript, and gene IDs, further complicates inter-database mapping, particularly when using versioned IDs. While G6PD has only two stable gene IDs, it has seven stable transcript IDs and thirteen versioned transcript IDs (**Figure 3B**). To assess how pervasive multi-mapping is across the whole CpDAA dataset, we quantified the mean number of Ensembl IDs per UniProtKB IDs. We found that the mean number of Ensembl IDs per UniProtKB IDs was significantly increased in versioned IDs compared to stable IDs, both for Ensembl gene and transcript IDs (**Figure EV2**). We suspected that the presence or absence of isoforms might contribute to the observed multi-mapping. Therefore, we extended our analysis of multi-mapping trends to compare the average number of Ensembl IDs mapping to single isoform- (**Figure 3D**) or multi-isoform- (**Figure 3E**) associated UniProtKB IDs. Not surprisingly, we found that UniProtKB stable identifiers with multiple associated protein isoforms have higher average counts of Ensembl ID types per UniProtKB identifier when compared to UniProtKB IDs associated with only one protein isoform. In addition, single isoform UniProtKB IDs are more likely to cross-reference identical Ensembl proteins compared to multi-isoform UniProtKB IDs mapping with stable identifiers (**Figure EV3 and EV4 and Table S11a, S11b)**. Therefore, single isoform proteins and their corresponding UniProtKB IDs are more resistant to residue-level mis-annotation when using stable IDs from the Ensembl release-specific mapping files to annotate CpDAA coordinates.

Next, we sought to determine how the changes to protein sequences associated with stable IDs varied across releases. We expected that temporally close releases of Ensembl would share higher sequence similarity and be less prone to mismapping. We identified the top stable IDs with poor sequence alignment across gene, transcript, and protein IDs (**Table S12a)**. We then used the Hamming(Frederick, Sedlmeyer, and White 1993) and Levenshtein(Yujian and Bo 2007) normalized distance to score protein sequence similarity on a scale of 0 to 1, where 0 indicates that the sequence is identical to the UniProtKB stable ID sequence. We found that the sequences associated with the UniProtKB and the Ensembl stable IDs varied significantly depending on the Ensembl version (**Figure 3F and EV5**), with temporally close releases showing generally greater sequence similarity. 49 UniProtKB IDs had no canonical sequence equivalent in all five Ensembl releases analyzed, with 17 of these IDs having CpDAA index differences for all detected cysteine and/or lysine positions (**Table S12b**).

One last challenge we identified is that the origin of the cross-reference file (whether it was created by UniProtKB or by Ensembl) affected the outcome of our mapping procedures. We pooled stable and versioned identifiers across five Ensembl releases and found that 56.9% of all Ensembl entries were identical to the UniProtKB ID (**Table S11b, Figure EV3**). We suspected that the absence of versioned identifiers in our CpDAA proteomics data likely contributed substantially to the observed multi-mapping when using a cross-reference file generated by Ensembl. We then used a cross reference file from UniProtKB that displays the CCDS canonical isoform ID mapped to a stable Ensembl protein ID, to more accurately identify the best mapped Ensembl protein ID. This approach allowed for > 99% sequence match across the mapped IDs and substantially reduced effects of multimapping with Ensembl IDs (**Table S11a, Figure EV4**). Our study demonstrated that mapping between the residue- and nucleotide-level requires attention to details regarding database updates, multi-mapping, and cross-reference file sources.

### (4) Assessment of pathogenicity predictions for CpD cysteine and lysine codons, using residue-codon mapping

Our next objective was to apply residue-codon mapping to the prioritization of functional CpDAAs—defined here as amino acids identified in our chemoproteomics studies that are more likely to control protein interactions, stability, localization or cellular abundance. Cysteines and lysines are both highly conserved, with 97% (Miseta and Csutora 2000) and 80% (Hacker et al. 2017) median level conservation, respectively. Consequently, sequence motif conservation cannot distinguish between functional and non-functional residues within chemoproteomic datasets. To identify cysteine- and lysine-centric genetic features suitable for pathogenicity prioritization, we tailored our pipeline to reverse-translate CpD cysteine and lysine positions to genomic coordinates of codons and genomic-based functional annotations. Cysteines and lysines were required to have valid coordinates in GRCh37 and GRCh38 reference genome assemblies, as some functional genetic variant annotations are only available for a one reference genome. For all proteins within our CpDAA dataset, we also processed both detected and undetected cysteines and lysines (**Figure EV6**). We found that probe-labeled cysteines and lysines represent ∼15% of all cysteines (6,057 CpD Cys out of 40,107 total Cys) and ∼6% of all lysines (8,868 CpD Lys out of 149,520 total Lys) found in chemoproteomic-identified proteins (n=3,840 UniProtKB IDs successfully mapped) (**Figure 4A and 4B**).

**Figure 4.**
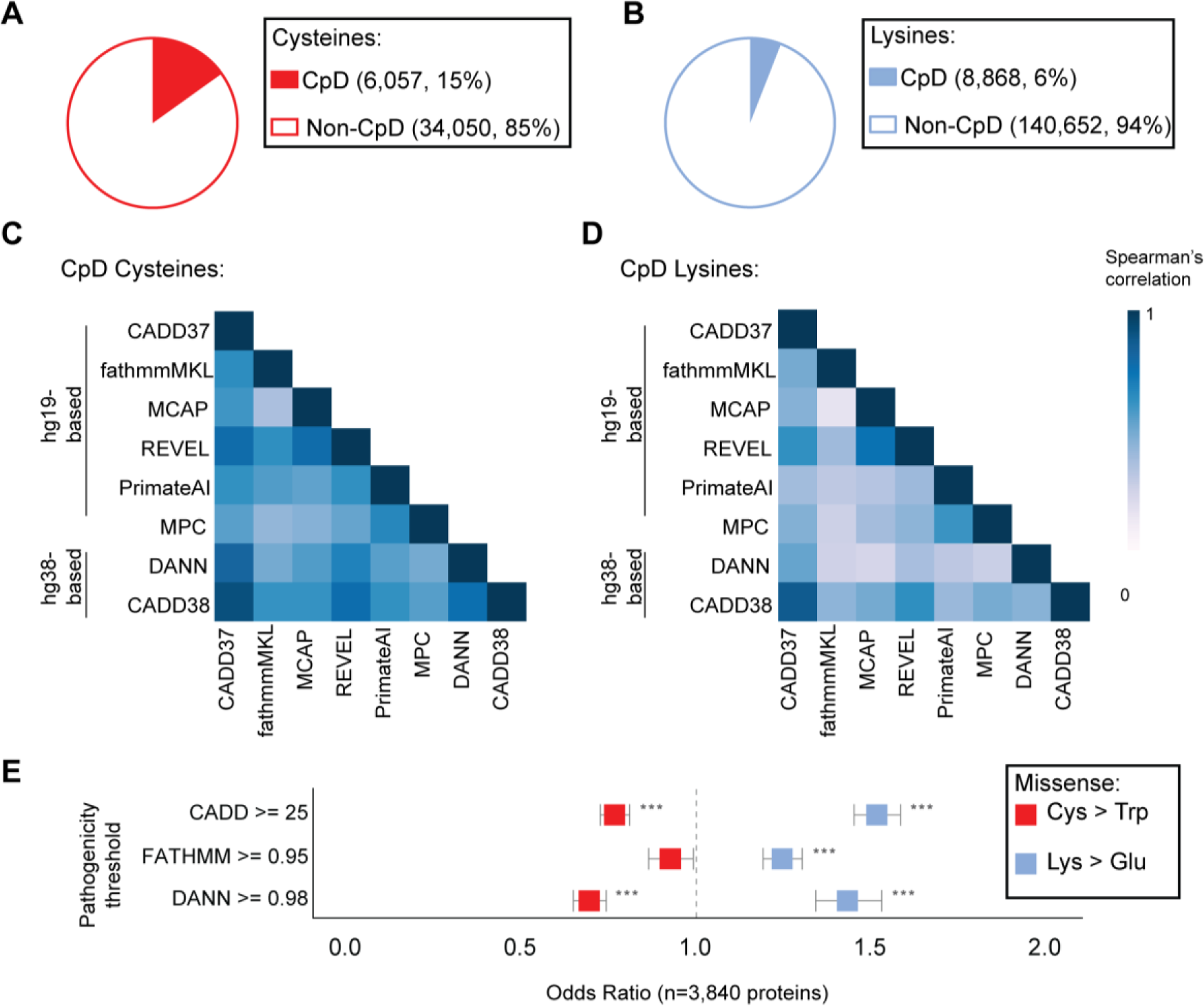
Mapping CpDAA genomic coordinates to predictions of pathogenicity. A-B) Aggregate number of cysteines (A) and lysines (B) in CpDAA-containing proteins (n=3,840), including both detected and undetected residues. C-D) Spearman’s correlation (r) of scores for all possible non-synonymous SNVs at cysteine (C) and Lysine (D) CpDAA codons. E) Odds ratio (OR) comparing the predicted pathogenicity of amino acid substitutions at detected versus undetected residue positions. Cys>Trp (yellow) and Lys>Glu (purple) missense scores were compared using a two-tailed Fisher’s Exact test with a 95% two sided confidence interval at the indicated score thresholds (y axis). OR> 1.0 indicates enrichment and OR< 1.0 indicates depletion of deleterious nonsynonymous SNVs at CpDAA relative to undetected residues. P-value cut-off = 5.0e-05, *** for values < 5.0e-21.

We next mapped the CpDAA coordinates to a panel of functional annotation scores (Kircher et al. 2014; Jagadeesh et al. 2016; Ioannidis et al. 2016; Rogers et al. 2018; Samocha et al. 2017; Quang, Chen, and Xie 2015; Sundaram et al. 2018), aiming to assess the concordance between individual scores and chemoproteomic identification. We selected complementary gene-, sub-gene-, and variant-level scores (**Table S13**) for our analysis. For variant-level pathogenicity predictors, higher scores correlate with increased importance of a residue to a protein’s function. To visualize the concordance of different scores for nonsynonymous SNVs, we calculated the correlation of scores for overlapping codon coordinates of CpD cysteine and lysine residues. While we observed generally high correlation for all scores, the correlation between pathogenicity predictions for CpD cysteine substitutions **(Figure 4B**) was higher (Spearman’s rank r between 0.36 and 0.91) than for CpD lysine substitutions (Spearman’s rank r between 0.16 and 0.81) (**Figure 4C**). When comparing only machine learning-based scores for substitutions at detected or undetected cysteines and lysines in chemoproteomic-detected proteins, we observed a lower correlation for detected compared to undetected lysines, and the opposite trend for cysteine residues (**Figure EV7**). We hypothesize that this higher correlation of annotation scores for cysteine may be driven in part by the high degree of cysteine conservation across evolution (Miseta and Csutora 2000) and the known critical functions of cysteines, including in catalysis, protein structure and redox regulation(Grunwell et al. 2015). For both CpD cysteines and CpD lysines, we observed a trend of slightly higher CADD scores with the GRCh38 model compared to GRCh37 model (**Table S14**).

Pathogenicity thresholds, which are provided by a subset of the scores investigated (e.g. CADD, FATHMM, and DANN), provide a useful cutoff for assessing whether substitutions at specific amino acids are likely to be deleterious to protein function. Therefore, we next assessed whether substitutions at detected versus undetected cysteines or lysines were more likely to be predicted damaging. Missense changes with high Grantham score (Grantham 1974), Cys>Trp and Lys>Glu, were used to assess the impact of these non-synonymous substitutions (missense) at detected versus undetected cysteine and lysine residues. We find that for CADDhg38 (Kircher et al. 2014) CADD GRCh38 model), FATHMM (Rogers et al. 2018), and DANN (Quang, Chen, and Xie 2015), mutations of detected cysteines are less likely to be predicted damaging compared to undetected cysteines **(Figure 4E, red**). In contrast, mutations of detected lysines were found to be more deleterious when compared to undetected lysines (**Figure 4E, blue**).

### (5) Chemoproteomic data combined with pathogenicity scores can help prioritize functional residues

We next assessed correlations between genetic-based pathogenicity score and amino acid reactivity, as assessed by chemoproteomics. We chose CADD as the optimal score to evaluate, as it integrates other nucleotide variant predictors into its model and is available for both reference genome assemblies, GRCh37 and GRCh38. Chemoproteomic reactivity measurements were binned into low, medium, and high reactivity categories, defined as low (R10:1>5), Medium (2<R10:1<5), High (R10:1<2) isoTOP-ABPP ratios, respectively(Weerapana et al. 2010; Hacker et al. 2017). These ratios quantify the relative labeling of a residue at different probe concentrations (e.g. 1× vs 10×). A ratio closer to one indicates that labeling is saturated at low probe concentration, which corresponds to a cysteine or lysine with higher intrinsic reactivity.

To adapt CADD scores from the nucleotide level to the amino acid level for CpDAAs, the mean and max CADD score for all possible nonsynonymous SNVs per codon (see Methods) were calculated. For both max (**Figure 5A)** and mean (**Figure EV8)** CADD codon scores, we found that highly reactive cysteines show significantly higher predicted deleteriousness (Cysteine CADD38 PHRED: Low 24.50 mean, 0.07 SD, 95% CI [24.35,24.64]; Medium 25.34 mean, 0.09 SD, 95% CI [25.16,25.53]; High 27.34 mean, 0.17 SD, 95% CI [27.01,27.66 CI]). In contrast, lysine reactivity did not correlate with predicted pathogenicity (**Figure 5B and Figure EV8**) (Lysine CADD38 PHRED: Low 26.12 mean, 0.02 SD, 95% CI [26.07,26.17]; Medium 25.90 mean, 0.06 SD, 95% CI [25.79,26.01]; High 26.07 mean, 0.08 SD, 95% CI [25.91,26.22]).

**Figure 5.**
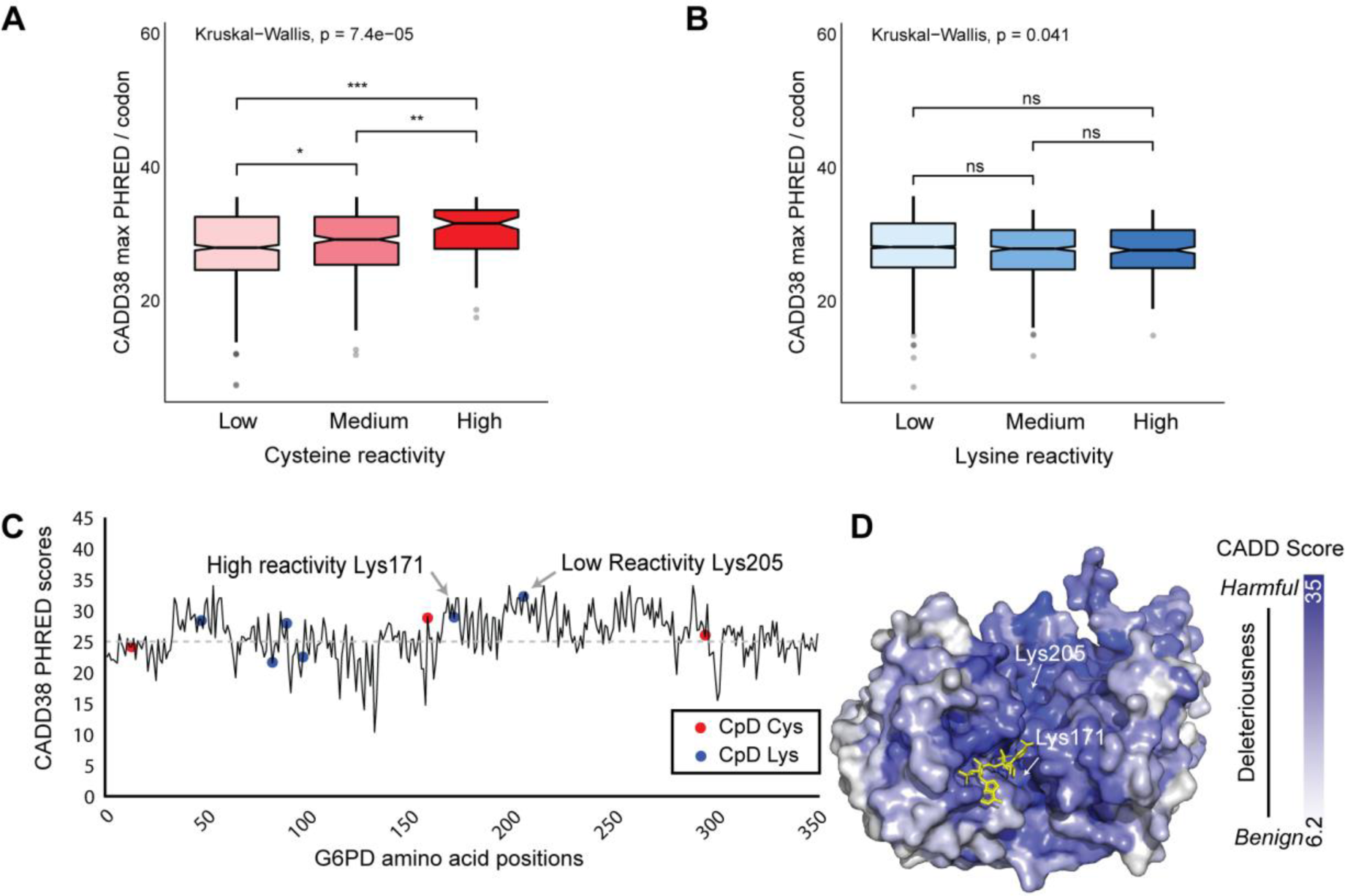
Association between amino acid reactivity and CADD score. A-B) Distribution of the max CADD38 (model for GRCh38) PHRED score/codon for (A) cysteine (n=1,401) and (B) lysine (n=4,363) CpDAAs of low, medium, and high intrinsic reactivities, defined by isoTOP-ABPP ratios, low (R10:1>5), Medium (2<R10:1<5), High (R<2) (Weerapana et al. 2010; Hacker et al. 2017). Kruskal-Wallis test was used for multiple pairwise-comparisons and Wilcox test was used for pairwise comparisons. Significant FDR adjusted p-values marked, *, *p* < 0.1, **, *p* < 0.01, ***, *p* < 0.0001. C) Shows CADD38 max codon missense scores for residues 1-300 of G6PD (UniProt ID P11413). D) Crystal structure of G6PD (PDB ID: 2BH9) shows K205 and K171 located within the enzyme active site. NADP^+^ cofactor shown in yellow. Surface colored by CADD38 max codon missense scores. Image generated in PyMOL (R. H. B. Smith, Dar, and Schlessinger, n.d.; DeLano and Others 2002).

As a first use-case to test the utility of integrating genetic-based pathogenicity predictions with CpDAA reactivity measures, we turned to the well characterized enzyme glucose-6-phosphate dehydrogenase (G6PD), which is an essential enzyme that when mutated results in one of the most common genetic enzymopathy caused by over 160 different point mutations (Hwang et al. 2018). G6PD deficiency is associated with both acute and chronic hemolytic anemia (Miwa and Fujii 1996; Porter et al. 1964) (OMIM #300908), and associated with malaria resistance (Luzzatto, Usanga, and Reddy 1969) (OMIM #611162). The G6PD protein contains 15 residues that were identified by chemoproteomics, 10 lysines and 5 cysteines. Of the CpD positions, K171 is the only highly reactive residue, and positions C13, K205, C385, K408, C446, and K497 are lowly reactive. Several residues, including K386 and C294 were not identified by our reactivity studies but were labeled in our studies aimed at identifying small molecule ligands(Backus et al. 2016; Hacker et al. 2017). Mutations at or affecting K171(Au et al. 2000) and K386 are implicated in anemia (Hirono et al. 1989).

To visualize CADD scores along protein sequence length, we plotted the first 300 amino acids in G6PD with lines tracking max CADD GRCh38 scores (**Figure 5C**). While K171 and K205 had very different intrinsic reactivities (high vs low, respectively), both showed high max CADD scores (28.8 and 32, respectively). Compared to other lowly reactive lysines in G6PD, lowly reactive K205 is the only residue with a missense average CADD score above the standard threshold of 25 for deleteriousness (K205 mean 26.3 compared to K408 mean 23.8 and K497 mean 23.3). Analysis of the molecular structure of G6PD (**Figure 5D** PDB ID: 2bh9) revealed that K171 and K205 are located in the enzyme active site, proximal to the nicotinamide-adenine-dinucleotide phosphate (NADP^+^) cofactor. Taken together, this analysis highlights the utility of integration of pathogenicity predictions to improve stratification of chemoproteomics data.

### (6) DISCUSSION

We conducted an in-depth analysis of multiple mapping strategies to facilitate multi-omic analysis of chemoproteomics datasets. We then applied our optimal mapping strategy to analyze how genetic variant predicted pathogenicity scores correlate with CpDAA data of measured intrinsic reactivity of cysteine and lysine residues. Our study revealed a number of challenges that limit the precision of multi-omic, residue-level data integration. Databases of genomic, transcriptomic and proteomic data adhere to different update cycles (Figure 1C), which results in different mapping outcomes depending on which version of the proteome, transcript and genome are used to process the raw proteomic data. These version updates impact both identifier mapping and mapping to residues and genomic coordinates.

Identification of sources of mismapping is an essential step to ensure high-quality and reproducible data integration. Proteomics research laboratories routinely share raw proteomic data files along with publication through the the ProteomeXchange Consortium (Deutsch et al. 2017) using resources such as PRIDE (Côté et al. 2012), PeptideAtlas, Massive (Deutsch et al. 2017), Panorama (Sharma et al. 2014), jPost (Rigden and Fernández 2019), and iProx (Rigden and Fernández 2019). However, providing the reference databases, which are typically a custom UniProtKB FASTA file, is not routine, making residue-level multi-omics integration challenging. Although UniProtKB is updated monthly, only annual releases are maintained long-term, meaning that the exact reference proteome data set used may no longer be publicly available for subsequent follow-up studies.

The availability of raw proteomic data might suggest an obvious solution: to <u>re-search raw</u> data using a new UniProtKB reference. However, reprocessing raw data can be both computationally expensive and time limiting. An important alternative is to <u>re-map</u> the processed data files to a newer and/or common UniProtKB reference to facilitate comparisons between datasets. However, in practice this approach can result in data loss and errors, which may confound interpretation. For example, when we re-mapped our legacy data to multiple UniProtKB version releases, we lost 28-199 of our proteins, which ranges from 0.6-4.8% of the original data (**Figure 1B**). While this may, at first glance seem to be a paltry fraction of all data, these losses can still prove problematic when key proteins of interest (**Table S7**) are lost due to mismapping. Making reference FASTA files publicly available alongside raw data files is a relatively simple solution to facilitate data integration.

There are several interconnected causes for our observed data loss at the protein level. The absence of protein isoform identifiers in most proteomics reference datasets, particularly when combined with database updates to canonical sequences can lead to mismapping, as shown for PRMT1 and FKBP7 (**Figure 2A**). The small number of UniProtKB sequences for which the canonical sequence is not the UniProtKB “-1” entry can also lead to further mismapping (**Figure 2C**). For protein-level mapping, the use of highly curated datasets, such as the Swiss-Prot reviewed CCDS-UniProtKB subset, can in part mitigate these losses.

Reversing the central dogma to map protein identifiers back to transcript and gene identifiers and amino acids back to genomic coordinates adds several additional layers of mapping complexity. Ensembl stable identifiers (gene, transcript, and protein), which are linked to UniProtKB stable identifiers are useful for facilitating this process. However, the number of redundant sequences maintained by Ensembl and the dynamic landscape of Ensembl entries across releases, complicates the use of Ensembl stable IDs for inter-database mapping. For example, for the protein G6PD, across the five Ensembl releases investigated, we identified seven stable protein IDs, of which only 1 was consistently identical to the UniProtKB canonical sequence for G6PD (**Figure 3A**). Database updates that result in changes to protein sequences associated with stable IDs are a particularly problematic cause of mismapping, particularly when assessed at the chemoproteomic residue-level. Across a number of commonly studied proteins, including MCM3, RAD50, and DIABLO, we found substantial differences in Ensembl protein and UniProtKB sequences across Ensembl releases (**Figure 3F**). Practically speaking, what this means is that a residue from a proteomics dataset could easily be mapped to the incorrect amino acid in an Ensembl protein, followed by the incorrect transcript position, incorrect genomic coordinates, and incorrect pathogenicity score.

Choice of reference genome further complicates data mapping. While many studies have now transitioned to GRCh38, many useful tools, including many predictions of pathogenicity (e.g. MPC, PrimaryAI, M-CAP, fathmm-MKL), were built using GRCh37. As GRCh37 was frozen in 2014, substantial mismapping is likely to occur when proteomics datasets generated using newer reference proteomes are mapped back to GRCh37. In the genetics community, this problem is commonly circumvented by performing a ‘lift-over’ from GRCh37 to GRCh38, bringing along the annotations linked with a specific reference genome. However, not all tools are compatible with liftover, as shown in **Table S13**. Local sequence alignment tools can be used to address the problems when transitioning between GRCh37 and GRCh38 but can be challenging to scale genome-wide. Transitioning all relevant annotations to the GRCh38 reference genome is ongoing and will address many of the aforementioned issues. However, this move is a substantial undertaking that requires rerunning of large-scale data sets and quality control.

Together our analysis of inter-database mapping enabled us to compile a rigorously curated dataset of CpDAAs that mapped to both GRCh37 and GRCh38 scores. Using this dataset, we were then able to ask a number of novel questions, including how different scores compare across all identified cysteine and lysine residues and whether the codons of specific residues are enriched for predicted pathogenic mutations. For all nucleotide substitutions that result in a cysteine or lysine amino acid change, we observed generally high concordance between scores (**Figure 4C and 4D**). While mutations at cysteine codons were observed to be, in general, predicted less deleterious than those at lysine codons, the subset of cysteines with heightened reactivity were predicted to be more damaging than cysteines of lower reactivity (**Figure 4E, 5A, EV8A**). No such trend was observed for highly reactive lysines (**Figure 4E, 5B, EV8B**). These intriguing findings suggest that cysteine hyperreactivity is a privileged feature that could be used to inform the functions of genetic variants.

We can foresee a multitude of applications for chemoproteomic and genomic data integration. While prior studies that revealed hyper-reactive cysteine and lysine residues are enriched in redox-active sites and enzyme active sites(Backus et al. 2016; Weerapana et al. 2010) and hyper-reactive lysines were depleted in post-translational modification sites (PTMs)(Hacker et al. 2017), most CpDAAs still lack functional annotation. Predictive tools, such as those highlighted here, will undoubtedly aid in stratification of residues identified by chemoproteomics studies, pinpointing potentially druggable and disease-linked pockets in proteins.

Another area that will undoubtedly benefit from such multi-omic approaches is interpretation of the impact of rare missense variation identified in patients with monogenic disorders. Protein-level functional data can aid in the interpretation of variants of uncertain significance (VUS), including those identified in clinical genetic testing, and can guide follow-up research studies. Proteomic approaches are also arguably more high-throughput relative genetic approaches, including site-directed mutagenesis. Application of chemoproteomics data to clinical studies will require careful data integration and sequence level mapping, particularly given that the reference sequences and choice of identifiers (versioned vs stable) employed by clinical vs. research studies are typically non-identical.

Addition of protein structural data to these pipelines will further improve their utility and predictive power. As a starting point to such structure-based data integration, we mapped CADD predictive scores directly to the structure of G6PD (**Figure 5D**). This 3-dimensional data integration highlighted key residues that form a common function in 3D space but are not easily identified using predictions associated with conservation in the linear-space of DNA. Looking to the future, we anticipate that such multi-omic studies will likely prove most enabling when combined with rigorous functional validation, for example by combining CRISPR-Cas9 mutagenesis with phenotypic assays. The use of CRISPR-Cas9 base editors (Kim et al. 2019; Grünewald et al. 2020, 2019) should facilitate such studies, particularly when combined with protein-centric guide RNA design packages (e.g. CRISPR-TAPE)(Anderson et al. 2020). In sum, we anticipate that such studies represent the next frontier for both the genetics and chemoproteomics communities.

## METHODS

### Data sources

Amino acid coordinates in this paper refer to UniProtKB-SwissProt human proteome filtered by canonical isoform and cross-reference in CCDS database downloaded August 06 2018 (2018_06) from the website (see URLs). These UniProtKB protein IDs were mapped to Ensembl IDs using two sources of cross-reference files, UniProtKB ID-mapping (idmapping.dat) or Ensembl release-specific mapping files (xref files) (Aken et al. 2016). UniProtKB sequences were aligned to ID cross-referenced Ensembl peptide sequences from releases v85, v92, v94, v96, and v97 FASTA files downloaded from the website (see URLs) November 19 2019. CADDv1.4 (Kircher et al. 2014) scores were downloaded from the website on July 03 2019. DANN (Quang, Chen, and Xie 2015), fathmmMKL (Shihab et al. 2014), M-CAP (Jagadeesh et al. 2016), MPC (Samocha et al. 2017), REVEL (Ioannidis et al. 2016), and PrimateAI (Sundaram et al. 2018) scores were obtained from dbNSFPv4.0a (Liu et al. 2016) downloaded from the website (see URLs) June 11 2019. Additional gene constraint score LOEUF from gnomADv2.1.1 (Karczewski et al. 2020) was downloaded March 22 2019.

https://www.uniprot.org/downloads

http://uswest.ensembl.org/info/data/ftp/index.html

https://cadd.gs.washington.edu/download

https://sites.google.com/site/jpopgen/dbNSFP

https://gnomad.broadinstitute.org/downloads

### Database update cycles

Average time between Ensembl, GENCODE, CCDS, and NCBI updates was quantified using all releases between August 2013 - July 2019 (5 years and 11 months window of time). Dates counted refer to the public release date posted on each databases’ ftp site. A total of 25 Ensembl, 13 GENCODE, 6 CCDS (only *homo sapien* releases), and 5 NCBI releases were used to calculate average time between each update. For UniProtKB update cycle length, values provided from the UniProtKB website on typical time between Knowledgebase releases from 2019 (4 weeks) and 2020 (8 weeks) were averaged. The timeline displays a subset of Ensembl, GENCODE, CCDS, and NCBI release dates used to calculate average database update cycles. Release dates were selected based on proximity to the dates of the five Ensembl releases analyzed for identifier multi-mapping.

### Mapping CpDAA data to more recent UniProtKB releases

Chemoproteomic-detected amino acid datasets had been previously searched against a non-redundant reverse concatenated UniProtKB reference FASTA file (Hacker et al. 2017; Backus et al. 2016; Weerapana et al. 2010) from the November 2012 (2012_11) release and amino acids in labeled peptides were annotated with the corresponding UniProt stable ID, amino acid letter, and position (e.g. P11413_C205). The author-provided UniProtKB 2012_11 FASTA file was referenced to check the UniProtKB IDs and CpDAA coordinates. Legacy chemoproteomic-detected cysteine and lysine residues that did not match amino acid letter and coordinate in the 2012_11 version of peptide sequence were removed prior to further analysis. The 2012-based CpDAA UniProtKB IDs and coordinates were then mapped to canonical sequences from the 2018_06 UniProtKB human proteome dataset using a custom python script.

### Criteria for removal from further analysis

Chemoproteomic-detected proteins were excluded from further analysis if: (1) UniProt canonical sequence from 2018 release was missing chemoproteomic-detected positions (example: natural variant overlaps detected cysteine position), (2) UniProt ID flagged with ‘caution’ on UniProt’s website (example: https://www.uniprot.org/uniprot/Q8WUH1), and (3) matching stable UniProt ID was not found in one of the five Ensembl release-specific mapping files downloaded.

### Assessment of isoforms per stable UniProtKB ID

The UniProtKB *homo sapien* FASTA file containing canonical and isoform sequences was downloaded August 06 2018. Isoform counts per UniProt entry were counted based on versioned IDs included in the FASTA file, excluding the canonical isoform, which is marked by the ID with no isoform name detail (e.g. P11413).

### Identification of UniProtKB canonical isoform ID numbers

UniProtKB canonical isoform ID numbers (e.g. P11413-X, ‘X’ representing the isoform name), were identified for only multi-isoform UniProtKB entries. The isoform name for the canonical sequence was identified by comparison of isoform-specific IDs in the FASTA file (used to count isoforms per stable UniProtKB ID) and the ID mapping (idat) file from UniProtKB August 01, 2018 release downloaded from the website August 06, 2018. The FASTA file marks the canonical ID with no isoform name, but the idat file displays the canonical ID isoform name. A custom python script compared all isoform-specific IDs for a single stable entry between the file types and annotated the isoform name associated with the canonical protein sequence in the FASTA file.

### Interdatabase mapping of CpDAA residues between UniProtKB and Ensembl proteins

Two methods were used to cross-reference equivalent Stable or Versioned protein IDs between UniProtKB and five Ensembl releases:

A. Ensembl isoform-less mapping: Ensembl xref files specific to each release studied (v85, v92, v94, v96, v97) were used for inter-database identifier mapping. All Ensembl gene, transcript, and protein stable IDs associated with 3,953 CpDAA stable UniProtKB IDs were pooled. Multi-mapping and sequence identity features between Ensembl proteins mapped to UniProtKB proteins grouped by single or multi-isoform entry status.
B. UniProt isoform-specific mapping: UniProtKB ID-mapping (idmapping.dat) file from August 01, 2018 release was used for inter-database identifier mapping. Isoform-specific UniProt IDs for multi-isoform entries and stable IDs for single isoform entries were used to cross-reference Ensembl stable protein IDs in the idat file. Similar to Method A, multi-mapping and sequence identity were assessed for mapped UniProt and Ensembl proteins.

### Assessing identifier multi-mapping between UniProtKB and Ensembl

From **Method A** ID mapping, all Ensembl IDs mapped to a Stable CpDAA UniProt ID were pooled from five Ensembl xref files. The mean number of unique Ensembl IDs (Versioned and Stable) per UniProt ID was then calculated both for UniProt entries with a single isoform and with multiple isoforms. Sequence equivalence between Ensembl and UniProtKB was assessed as a boolean (True identical sequence or False for not identical) using a custom python script. From **Method B** ID mapping, stable Ensembl protein IDs pulled from the UniProtKB idat file were used to filter Ensembl release-specific FASTA files for associated sequences. Similar to analysis from **Method A**, multi-mapping was calculated both for single and multi-isoform UniProt entries with sequence equivalence determined using a custom python script. Statistical significance for all multi-mapping results was assessed with a Student’s unpaired T-test.

### Distances between mapped UniProtKB and Ensembl sequences across releases

ID mappings were obtained from **Method B**, using isoform-specific information to map UniProtKB canonical IDs for multiple isoform entries to equivalent stable Ensembl protein IDs. Protein sequence similarity was scored for mapped IDs using the Hamming normalized distance metric (Frederick, Sedlmeyer, and White 1993) and the Levenshtein normalized distance metric (Yujian and Bo 2007) to quantify how different the sequence is compared to the UniProtKB Stable ID sequence. Normalized scale is over 0 to 1, with 0 meaning the Ensembl protein is identical to the UniProt canonical sequence while 1 shows significant differences between the two sequences.

### Identification of frequently updated Ensembl gene, transcript, and protein IDs

Ensembl Stable IDs found in all five Ensembl releases studied were first identified. In total, 8,861 unique Ensembl protein IDs with associated transcript and gene IDs were used to create Stable Key IDs (formatted as ‘ENSG_ENST_ENSP’). These Ensembl key IDs cross-referenced 3,887 unique CpDAA UniProtKB IDs, and were used to filter Ensembl release-specific FASTA files for associated gene, transcript, and protein versioned IDs. To identify updating Ensembl sequences over the five studied releases, version number increments (signifying sequence re-annotation updates) since the v85 release were summed for a given gene, transcript, and protein ID (e.g. ID extension number ‘.X’). For comparison of gene, transcript, and protein ID types, version number sums for each release were calculated and compared to the v85 sum. To identify ‘dated’ ID mappings in which the Ensembl proteins are no longer equivalent to the canonical UniProtKB protein from 2018_06 release, we used ID cross-references from **Method B** and scored sequence distance by Hamming or Levenshtein metrics. We included 75 Ensembl Stable IDs with mapped proteins equivalent to the UniProtKB canonical sequence in v92 but significantly different in other Ensembl releases analyzed.

### Residue mapping to pathogenicity scores

CpDAA coordinates were mapped to dbNSFPv4.0. Matching UniProt Stable ID, and amino acid positions matching our CpDAA dataset were required. Coordinates for undetected cysteine and lysine residues were also pulled with dbNSFPv4.0. Reference genome GRCh37 and GRCh38 coordinates from dbNSFPv4.0 were then used to map CADDv1.4 scores from both CADD models (GRCh37 and GRCh38 models). Cysteines and lysines with valid coordinates in both assemblies were retained. All possible non-synonymous variant scores were required to be present (both for detected and undetected residues). Codons lacking variant scores were excluded from analysis. Both the max CADD score at a detected residue or the mean of all non-synonymous variants for that residue’s codon coordinates were assessed.

### Score thresholds for pathogenicity

CADD score greater than or equal to 25, fathmm-mkl score greater than or equal to 0.95, DANN score greater than or equal to 0.98.

### Mapping CADD scores to PDB structures

Max and mean non-synonymous CADD scores for all amino acid positions of protein G6PD (UniProtKB ID P11413) were calculated using downloaded CADD files (see URLs) and custom python script. Max and mean codon scores for the canonical sequence were next mapped to all amino acid positions found in the protein structure of G6PD (PDB: 2BH9). Beta factor values for alpha carbons of the structure were set to reflect CADD max or mean scores calculated.

### Data and source code availability

Analyses utilized Python 3.7.4 and R 3.6.2. Plots were generated in R using ggplot2 (Wickham 2016), Prism 8 (GraphPad Software Inc.), or Illustrator (Adobe Inc.). Code is available at https://github.com/mfpfox/MAPPING

## Supporting information

Expanded View Figures

## Abbreviations

(UniProtKB-SP): <u>U</u>niProt <u>K</u>nowledge <u>B</u>ase-<u>S</u>wiss-<u>P</u>rot,
(xref): E<u>x</u>ternal <u>Ref</u>erence,
(ENSP): <u>Ens</u>embl <u>P</u>rotein,
(ENST): <u>Ens</u>embl <u>T</u>ranscript,
(ENSG): <u>Ens</u>embl <u>G</u>ene,
(UKB): <u>U</u>niProt <u>K</u>nowledge <u>B</u>ase,
(CADD): Combined Annotation Dependent Depletion,
(CpDAA): <u>C</u>hemo<u>p</u>roteomic <u>D</u>etected <u>A</u>mino <u>A</u>cids,
(CCDS): <u>C</u>onsensus <u>C</u>o<u>d</u>ing <u>S</u>equence,
(DANN): <u>D</u>eleterious <u>A</u>nnotation of genetic variants using <u>N</u>eural <u>N</u>etworks,
(FATHMM-MKL): <u>F</u>unctional <u>A</u>nalysis through <u>H</u>idden <u>M</u>arkov <u>M</u>odels,
(PDB): <u>P</u>rotein <u>D</u>ata <u>B</u>ank,
(SNV): single nucleotide variant,
(dbNSFP): <u>D</u>ata<u>b</u>ase for <u>N</u>on <u>S</u>ynonymous <u>F</u>unctional <u>p</u>redictions.

## Data and source code availability

Analyses utilized Python 3.7.4 and R 3.6.2. Data and code available at the github site below are sufficient to reproduce the plots and analyses in this paper are available at https://github.com/mfpfox/MAPPING

## Software and Websites Accessed

https://www.uniprot.org/downloads

http://uswest.ensembl.org/info/data/ftp/index.html

https://cadd.gs.washington.edu/download

https://sites.google.com/site/jpopgen/dbNSFP

## Author Contributions

MFP, VAA, and KMB conceived of and designed the study. MFP wrote code, analyzed data sets and made figures. MFP, VAA, and KMB interpreted the results and wrote the manuscript.

## Notes

The authors declare no competing interests.

## Acknowledgements

This study was supported by a Beckman Young Investigator Award (Backus), V Scholar Award V2019-017 (Backus), Chemistry Biology Interface Training Program (T32GM008496 to Palafox) and DP5OD024579 (Arboleda). We gratefully acknowledge all members of the Backus and Arboleda labs for their helpful suggestions.

## References

Agrawal, Raag, and Sudhakaran Prabakaran. 2020. “Big Data in Digital Healthcare: Lessons Learnt and Recommendations for General Practice.” Heredity 124 (4): 525–34.

Aken, Bronwen L., Sarah Ayling, Daniel Barrell, Laura Clarke, Valery Curwen, Susan Fairley, Julio Fernandez Banet, et al. 2016. “The Ensembl Gene Annotation System.” Database: The Journal of Biological Databases and Curation 2016 (June). https://doi.org/10.1093/database/baw093.

Anderson, Daniel Paolo, Henry James Benns, Edward William Tate, and Matthew Andrew Child. 2020. “CRISPR-TAPE: Protein-Centric CRISPR Guide Design for Targeted Proteome Engineering.” Molecular Systems Biology 16 (6): e9475.

Au, S. W., S. Gover, V. M. Lam, and M. J. Adams. 2000. “Human Glucose-6-Phosphate Dehydrogenase: The Crystal Structure Reveals a Structural NADP(+) Molecule and Provides Insights into Enzyme Deficiency.” Structure 8 (3): 293–303.

Backus, K. M., B. E. Correia, K. M. Lum, S. Forli, B. D. Horning, G. E. Gonzalez-Paez, S. Chatterjee, et al. 2016. “Proteome-Wide Covalent Ligand Discovery in Native Biological Systems.” Nature 534 (7608): 570–74.

Côté, Richard G., Johannes Griss, José A. Dianes, Rui Wang, James C. Wright, Henk W. P. van den Toorn, Bas van Breukelen, et al. 2012. “The PRoteomics IDEntification (PRIDE) Converter 2 Framework: An Improved Suite of Tools to Facilitate Data Submission to the PRIDE Database and the ProteomeXchange Consortium.” Molecular & Cellular Proteomics: MCP 11 (12): 1682–89.

David, Fabrice P. A., and Yum L. Yip. 2008. “SSMap: A New UniProt-PDB Mapping Resource for the Curation of Structural-Related Information in the UniProt/Swiss-Prot Knowledgebase.” BMC Bioinformatics 9 (September): 391.

DeLano, Warren L., and Others. 2002. “Pymol: An Open-Source Molecular Graphics Tool.” CCP4 Newsletter on Protein Crystallography 40 (1): 82–92.

Deutsch, Eric W., Attila Csordas, Zhi Sun, Andrew Jarnuczak, Yasset Perez-Riverol, Tobias Ternent, David S. Campbell, et al. 2017. “The ProteomeXchange Consortium in 2017: Supporting the Cultural Change in Proteomics Public Data Deposition.” Nucleic Acids Research 45 (D1): D1100–1106.

Durinck, Steffen, Paul T. Spellman, Ewan Birney, and Wolfgang Huber. 2009. “Mapping Identifiers for the Integration of Genomic Datasets with the R/Bioconductor Package biomaRt.” Nature Protocols 4 (8): 1184–91.

“Ensembl Stable IDs.” n.d. Accessed July 1, 2020. https://uswest.ensembl.org/info/genome/stable_ids/index.html.

Frederick, William G., Robert L. Sedlmeyer, and Curt M. White. 1993. “The Hamming Metric in Genetic Algorithms and Its Application to Two Network Problems.” In Proceedings of the 1993 ACM/SIGAPP Symposium on Applied Computing: States of the Art and Practice, 126–30. SAC ‘93. New York, NY, USA: Association for Computing Machinery.

Gong, Sungsam, James S. Ware, Roddy Walsh, and Stuart A. Cook. 2014. “NECTAR: A Database of Codon-Centric Missense Variant Annotations.” Nucleic Acids Research 42 (Database issue): D1013–19.

Grantham, R. 1974. “Amino Acid Difference Formula to Help Explain Protein Evolution.” Science 185 (4154): 862–64.

Grünewald, Julian, Ronghao Zhou, Sowmya Iyer, Caleb A. Lareau, Sara P. Garcia, Martin J. Aryee, and J. Keith Joung. 2019. “CRISPR DNA Base Editors with Reduced RNA off-Target and Self-Editing Activities.” Nature Biotechnology 37 (9): 1041–48.

Grünewald, Julian, Ronghao Zhou, Caleb A. Lareau, Sara P. Garcia, Sowmya Iyer, Bret R. Miller, Lukas M. Langner, Jonathan Y. Hsu, Martin J. Aryee, and J. Keith Joung. 2020. “A Dual-Deaminase CRISPR Base Editor Enables Concurrent Adenine and Cytosine Editing.” Nature Biotechnology, June. https://doi.org/10.1038/s41587-020-0535-y.

Grunwell, Jocelyn R., Scott E. Gillespie, Janine M. Ward, Anne M. Fitzpatrick, Lou Ann Brown, Theresa W. Gauthier, and Kiran B. Hebbar. 2015. “Comparison of Glutathione, Cysteine, and Their Redox Potentials in the Plasma of Critically Ill and Healthy Children.” Frontiers in Pediatrics 3 (May): 46.

Guo, Yan, Yulin Dai, Hui Yu, Shilin Zhao, David C. Samuels, and Yu Shyr. 2017. “Improvements and Impacts of GRCh38 Human Reference on High Throughput Sequencing Data Analysis.” Genomics 109 (2): 83–90.

Hacker, S. M., K. M. Backus, M. R. Lazear, S. Forli, B. E. Correia, and B. F. Cravatt. 2017. “Global Profiling of Lysine Reactivity and Ligandability in the Human Proteome.” Nature Chemistry 9 (12): 1181–90.

Hirono, A., W. Kuhl, T. Gelbart, L. Forman, V. F. Fairbanks, and E. Beutler. 1989. “Identification of the Binding Domain for NADP+ of Human Glucose-6-Phosphate Dehydrogenase by Sequence Analysis of Mutants.” Proceedings of the National Academy of Sciences of the United States of America 86 (24): 10015–17.

Hwang, Sunhee, Karen Mruk, Simin Rahighi, Andrew G. Raub, Che-Hong Chen, Lisa E. Dorn, Naoki Horikoshi, Soichi Wakatsuki, James K. Chen, and Daria Mochly-Rosen. 2018. “Correcting Glucose-6-Phosphate Dehydrogenase Deficiency with a Small-Molecule Activator.” Nature Communications 9 (1): 4045.

Ioannidis, Nilah M., Joseph H. Rothstein, Vikas Pejaver, Sumit Middha, Shannon K. McDonnell, Saurabh Baheti, Anthony Musolf, et al. 2016. “REVEL: An Ensemble Method for Predicting the Pathogenicity of Rare Missense Variants.” American Journal of Human Genetics 99 (4): 877–85.

Jagadeesh, K. A., A. M. Wenger, M. J. Berger, H. Guturu, P. D. Stenson, D. N. Cooper, J. A. Bernstein, and G. Bejerano. 2016. “M-CAP Eliminates a Majority of Variants of Uncertain Significance in Clinical Exomes at High Sensitivity.” Nature Genetics 48 (12): 1581–86.

Karczewski, Konrad J., Laurent C. Francioli, Grace Tiao, Beryl B. Cummings, Jessica Alföldi, Qingbo Wang, Ryan L. Collins, et al. 2020. “The Mutational Constraint Spectrum Quantified from Variation in 141,456 Humans.” Nature 581 (7809): 434–43.

Kim, Daesik, Da-Eun Kim, Gyeorae Lee, Sung-Ik Cho, and Jin-Soo Kim. 2019. “Genome-Wide Target Specificity of CRISPR RNA-Guided Adenine Base Editors.” Nature Biotechnology 37 (4): 430–35.

Kircher, M., D. M. Witten, P. Jain, B. J. O’Roak, G. M. Cooper, and J. Shendure. 2014. “A General Framework for Estimating the Relative Pathogenicity of Human Genetic Variants.” Nature Genetics 46 (3): 310–15.

Liu, Xiaoming, Chunlei Wu, Chang Li, and Eric Boerwinkle. 2016. “dbNSFP v3.0: A One-Stop Database of Functional Predictions and Annotations for Human Nonsynonymous and Splice-Site SNVs.” Human Mutation 37 (3): 235–41.

Luzzatto, L., F. A. Usanga, and S. Reddy. 1969. “Glucose-6-Phosphate Dehydrogenase Deficient Red Cells: Resistance to Infection by Malarial Parasites.” Science 164 (3881): 839–42.

McLaren, William, Laurent Gil, Sarah E. Hunt, Harpreet Singh Riat, Graham R. S. Ritchie, Anja Thormann, Paul Flicek, and Fiona Cunningham. 2016. “The Ensembl Variant Effect Predictor.” Genome Biology 17 (1): 122.

Miseta, Attila, and Peter Csutora. 2000. “Relationship Between the Occurrence of Cysteine in Proteins and the Complexity of Organisms.” Molecular Biology and Evolution 17 (8): 1232–39.

Miwa, S., and H. Fujii. 1996. “Molecular Basis of Erythroenzymopathies Associated with Hereditary Hemolytic Anemia: Tabulation of Mutant Enzymes.” American Journal of Hematology 51 (2): 122–32.

Porter, I. H., S. H. Boyer, E. J. Watson-Williams, A. Adam, A. Szeinberg, and M. Siniscalco. 1964. “VARIATION OF GLUCOSE-6-PHOSPHATE DEHYDROGENASE IN DIFFERENT POPULATIONS.” The Lancet 1 (7339): 895–99.

Quang, Daniel, Yifei Chen, and Xiaohui Xie. 2015. “DANN: A Deep Learning Approach for Annotating the Pathogenicity of Genetic Variants.” Bioinformatics 31 (5): 761–63.

Rigden, Daniel J., and Xosé M. Fernández. 2019. “The 26th Annual Nucleic Acids Research Database Issue and Molecular Biology Database Collection.” Nucleic Acids Research 47 (D1): D1–7.

Rogers, Mark F., Hashem A. Shihab, Matthew Mort, David N. Cooper, Tom R. Gaunt, and Colin Campbell. 2018. “FATHMM-XF: Accurate Prediction of Pathogenic Point Mutations via Extended Features.” Bioinformatics 34 (3): 511–13.

Ruffier, Magali, Andreas Kähäri, Monika Komorowska, Stephen Keenan, Matthew Laird, Ian Longden, Glenn Proctor, et al. 2017. “Ensembl Core Software Resources: Storage and Programmatic Access for DNA Sequence and Genome Annotation.” Database: The Journal of Biological Databases and Curation 2017 (1). https://doi.org/10.1093/database/bax020.

Samocha, Kaitlin E., Jack A. Kosmicki, Konrad J. Karczewski, Anne H. O’Donnell-Luria, Emma Pierce-Hoffman, Daniel G. MacArthur, Benjamin M. Neale, and Mark J. Daly. 2017. “Regional Missense Constraint Improves Variant Deleteriousness Prediction.” bioRxiv. https://doi.org/10.1101/148353.

Schwartz, Gregory W., Tair Shauli, Michal Linial, and Uri Hershberg. 2019. “Serine Substitutions Are Linked to Codon Usage and Differ for Variable and Conserved Protein Regions.” Scientific Reports 9 (1): 17238.

Sehnal, David, Mandar Deshpande,Radka Svobodová Vařeková, Saqib Mir, Karel Berka, Adam Midlik, Lukáš Pravda, Sameer Velankar, and Jaroslav Koča. 2017. “LiteMol Suite: Interactive Web-Based Visualization of Large-Scale Macromolecular Structure Data.” Nature Methods 14 (12): 1121–22.

Sharma, Vagisha, Josh Eckels, Greg K. Taylor, Nicholas J. Shulman, Andrew B. Stergachis, Shannon A. Joyner, Ping Yan, et al. 2014. “Panorama: A Targeted Proteomics Knowledge Base.” Journal of Proteome Research 13 (9): 4205–10.

Shihab, Hashem A., Julian Gough, Matthew Mort, David N. Cooper, Ian N. M. Day, and Tom R. Gaunt. 2014. “Ranking Non-Synonymous Single Nucleotide Polymorphisms Based on Disease Concepts.” Human Genomics 8 (June): 11.

Sivley, R. Michael, Xiaoyi Dou, Jens Meiler, William S. Bush, and John A. Capra. 2018. “Comprehensive Analysis of Constraint on the Spatial Distribution of Missense Variants in Human Protein Structures.” American Journal of Human Genetics 102 (3): 415–26.

Smith, Lloyd M., Paul M. Thomas, Michael R. Shortreed, Leah V. Schaffer, Ryan T. Fellers, Richard D. LeDuc, Trisha Tucholski, et al. 2019. “A Five-Level Classification System for Proteoform Identifications.” Nature Methods 16 (10): 939–40.

Smith, Ryan H. B., Arvin C. Dar, and Avner Schlessinger. n.d. “PyVOL: A PyMOL Plugin for Visualization, Comparison, and Volume Calculation of Drug-Binding Sites.” https://doi.org/10.1101/816702.

Stephenson, James D., Roman A. Laskowski, Andrew Nightingale, Matthew E. Hurles, and Janet M. Thornton. 2019. “VarMap: A Web Tool for Mapping Genomic Coordinates to Protein Sequence and Structure and Retrieving Protein Structural Annotations.” Bioinformatics 35 (22): 4854–56.

Sundaram, Laksshman, Hong Gao, Samskruthi Reddy Padigepati, Jeremy F. McRae, Yanjun Li, Jack A. Kosmicki, Nondas Fritzilas, et al. 2018. “Predicting the Clinical Impact of Human Mutation with Deep Neural Networks.” Nature Genetics 50 (8): 1161–70.

Tang, J., A. Frankel, R. J. Cook, S. Kim, W. K. Paik, K. R. Williams, S. Clarke, and H. R. Herschman. 2000. “PRMT1 Is the Predominant Type I Protein Arginine Methyltransferase in Mammalian Cells.” The Journal of Biological Chemistry 275 (11): 7723–30.

Walkinshaw, Donald R., Ryan Weist, Go-Woon Kim, Linya You, Lin Xiao, Jianyun Nie, Cathy S. Li, Songping Zhao, Minghong Xu, and Xiang-Jiao Yang. 2013. “The Tumor Suppressor Kinase LKB1 Activates the Downstream Kinases SIK2 and SIK3 to Stimulate Nuclear Export of Class IIa Histone Deacetylases.” The Journal of Biological Chemistry 288 (13): 9345–62.

Weerapana, E., C. Wang, G. M. Simon, F. Richter, S. Khare, M. B. Dillon, D. A. Bachovchin, K. Mowen, D. Baker, and B. F. Cravatt. 2010. “Quantitative Reactivity Profiling Predicts Functional Cysteines in Proteomes.” Nature 468 (7325): 790–95.

Wickham, Hadley. 2016. “Programming with ggplot2.” In ggplot2: Elegant Graphics for Data Analysis, edited by Hadley Wickham, 241–53. Cham: Springer International Publishing.

Yujian, Li, and Liu Bo. 2007. “A Normalized Levenshtein Distance Metric.” IEEE Transactions on Pattern Analysis and Machine Intelligence 29 (6): 1091–95.

