## Supplementary material for "From Chemoproteomic-Detected Amino Acids to Genomic Coordinates: Insights into Precise Multi-omic Data Integration": Expanded View Figures

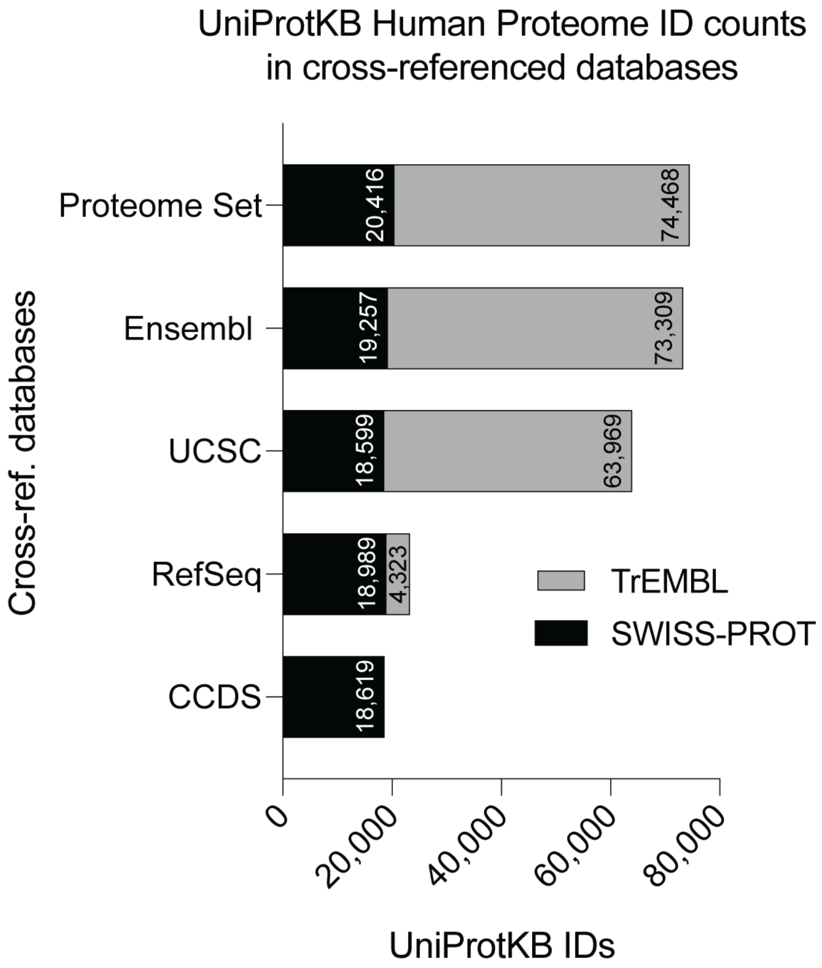


**Figure EV1**. UniprotKB Human Proteome ID counts in cross-referenced databases. The UniProtKB/TrEMBL subset (automated translations of coding sequences) are shown in grey and the UniProtKB/Swiss-Prot subset (manually curated sequences) are shown in black. Ensembl, USCS, and RefSeq contain both automated (TrEMBL) and manually curated (Swiss-Prot) entries. Sequences derived from the consensus coding sequence (CCDS) project are associated with the UniProtKB/Swiss-Prot subset.

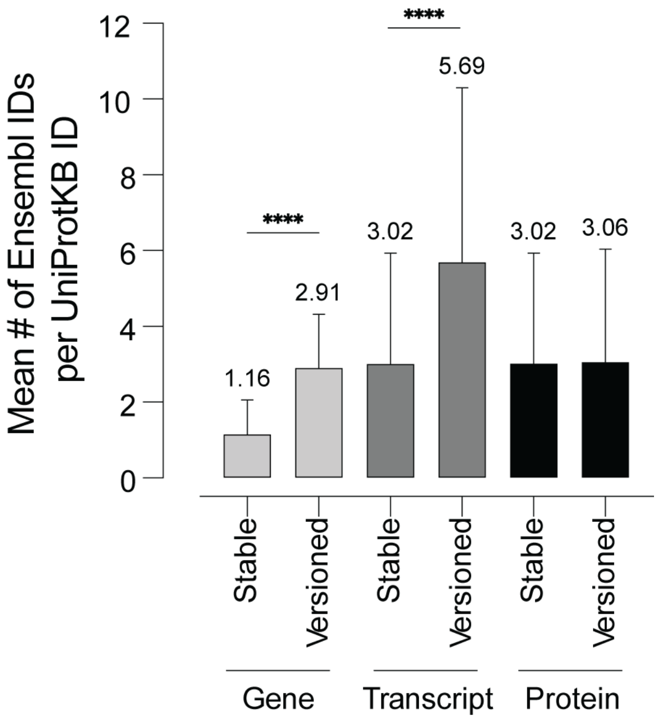


**Figure EV2.** Average number of Ensembl gene, transcript, and protein IDs for 3,953 CpD UniProtKB stable IDs. Multi-mapping was observed for stable (mean= 3.02 identifiers/UniProt ID) and versioned (mean= 5.69 identifiers/UniProtKB ID) Ensembl transcript IDs and for stable (mean= 3.02 identifiers/UniProt ID) and versioned (mean= 3.06 identifiers/UniProtKB ID) Ensembl protein IDs. Bar plots represent mean values ± SD for the number of Ensembl IDs per stable UniProtKB ID. Statistical significance was calculated using an unpaired Student's T-test, **** p-value <0.0001.

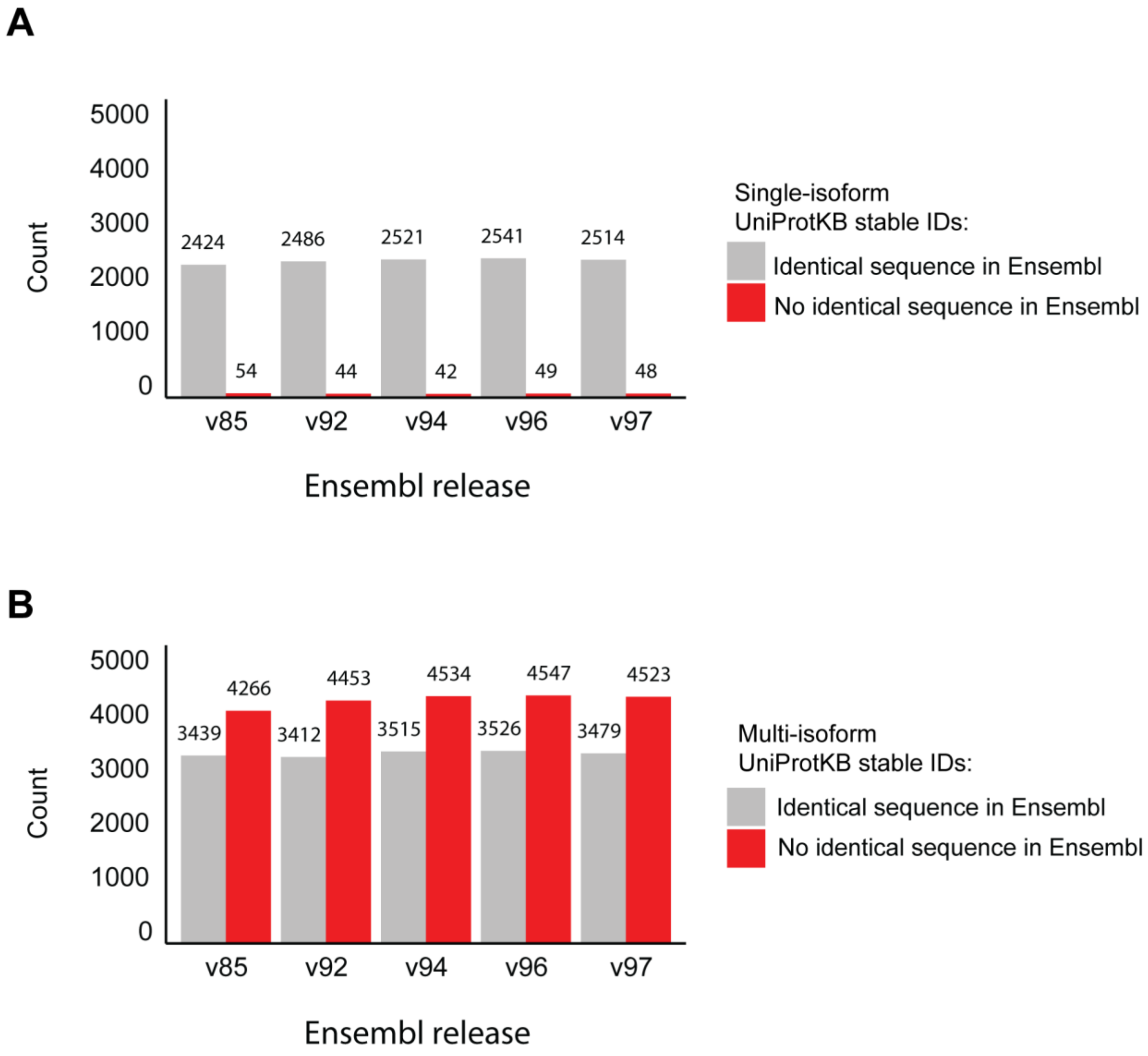


**Figure EV3.** Using Ensembl xref files, Ensembl protein IDs from five Ensembl releases were mapped to A) 1,466 single isoform UniProKB IDs and B) 2,487 multi-isoform UniProKB IDs.

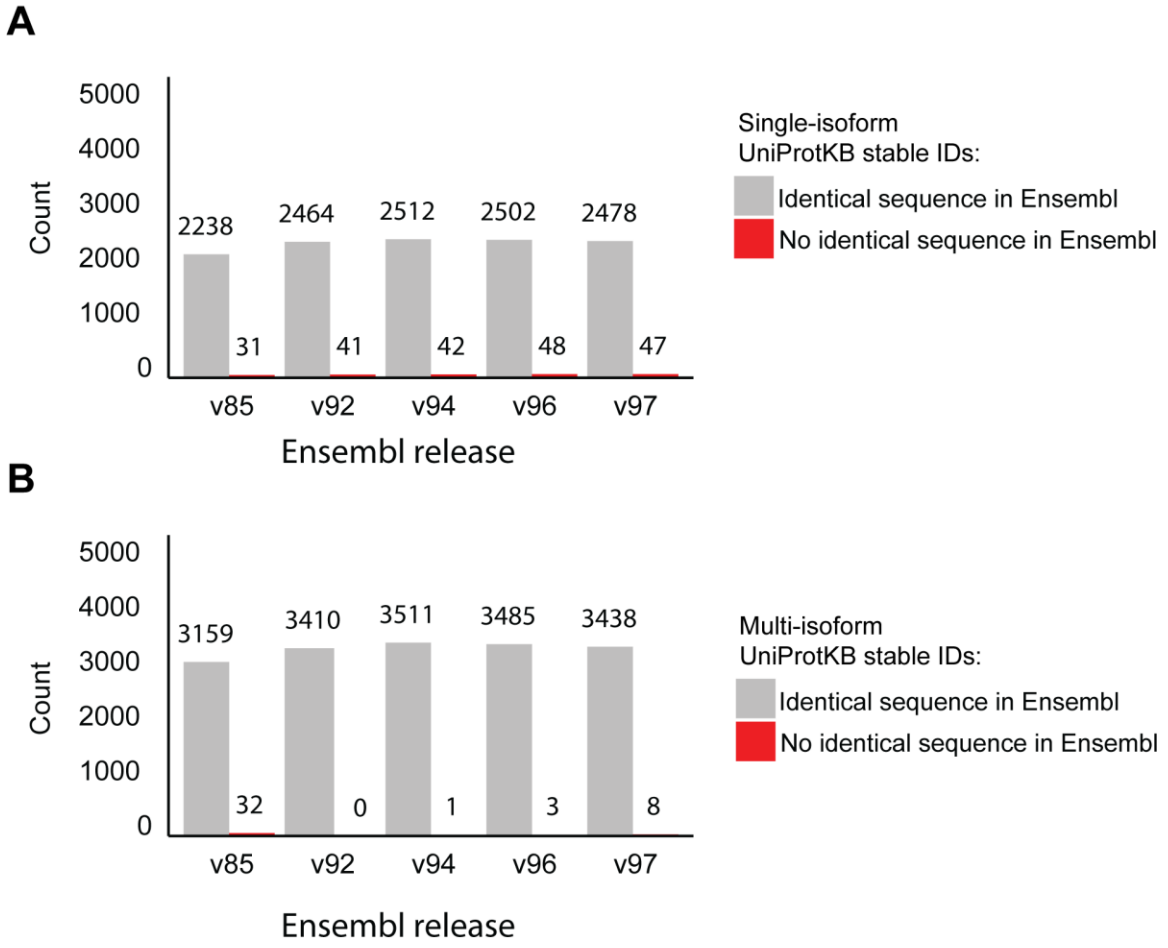


**Figure EV4.** Using UniProtKB provided versioned ID cross-reference files, Ensembl protein IDs from five Ensembl releases were mapped to A) 1,466 single isoform UniProtKB IDs and B) 2,487 multi-isoform UniProKB IDs.

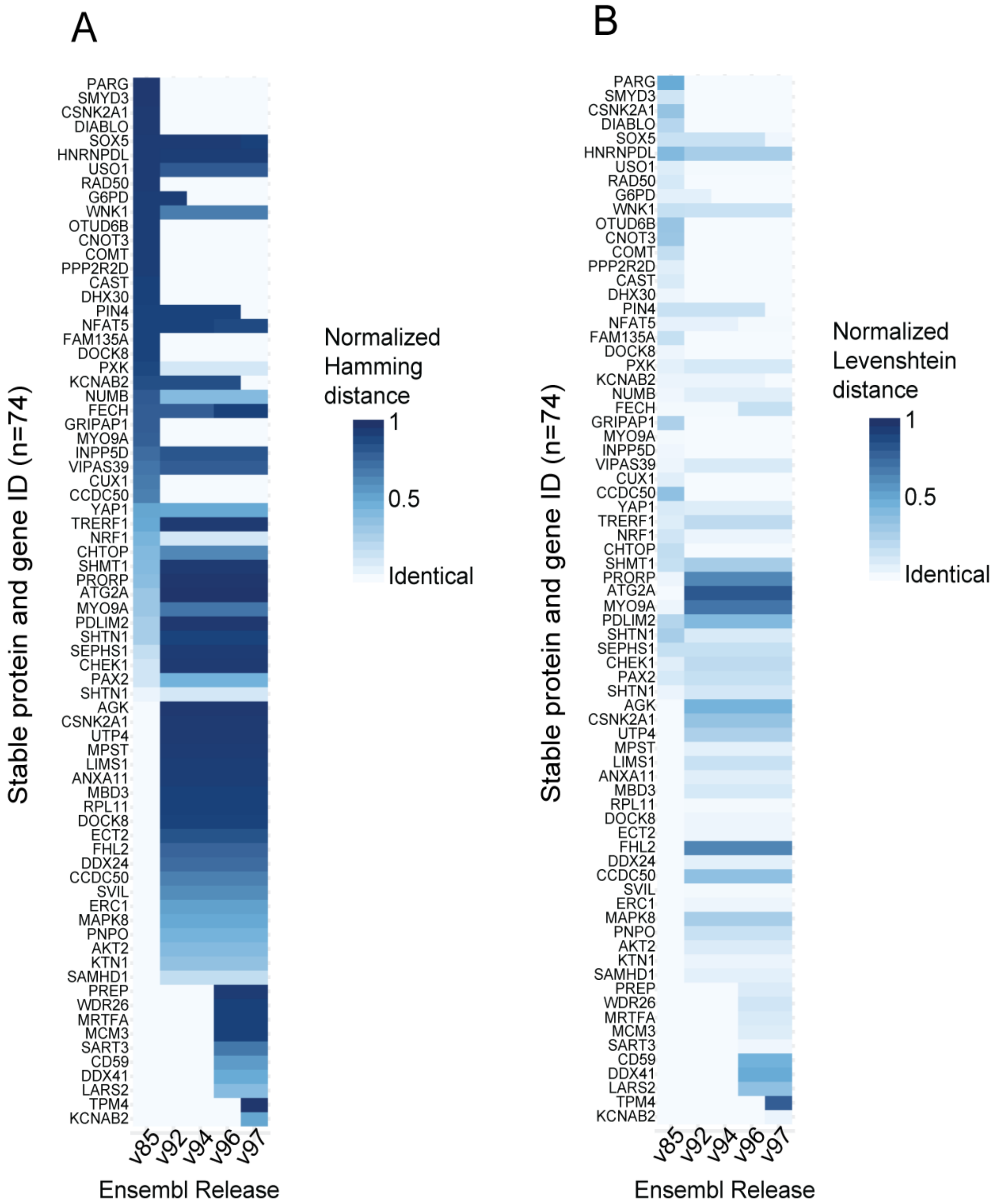


**Figure EV5.** Sequence similarity between UniProtKB protein sequences and protein sequences associated with Ensembl stable IDs across releases. Heatmaps show A) normalized Hamming distance and B) normalized Levenshtein distance for sequence alignments of the protein sequences associated with the top 74 stable Ensembl gene, transcript, and protein IDs with poor sequence alignments to the cross-referenced stable UniProtKB IDs across the indicated releases. Scores range from 0 to 1, with 0 indicating identical to the canonical sequence in the 2018 UniProtKB CCDS release.

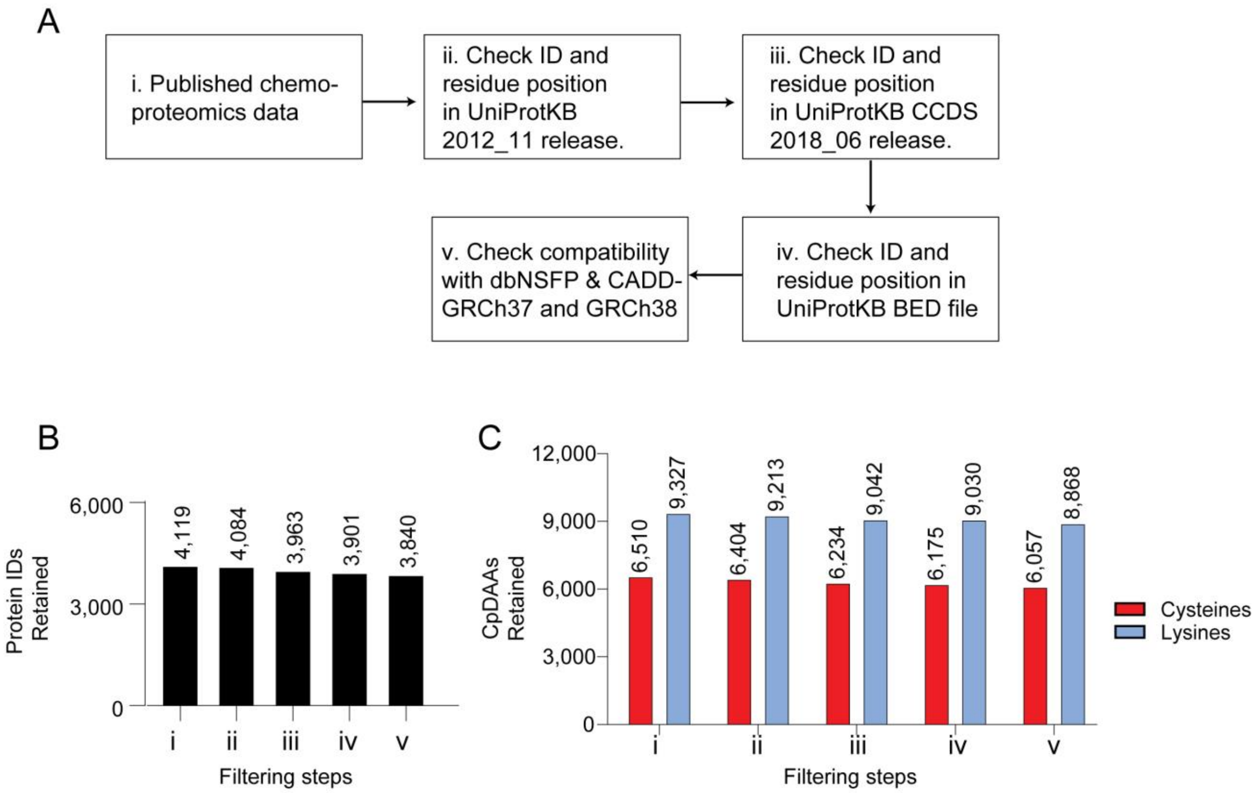


**Figure EV6.** Data retention during residue-codon mapping to pathogenicity scores. A) Shows workflow. B) Number of IDs retained after each mapping step shown in ‘A.’ C) Number of CpD cysteine and lysine residues retained during each mapping step shown in ‘A.’

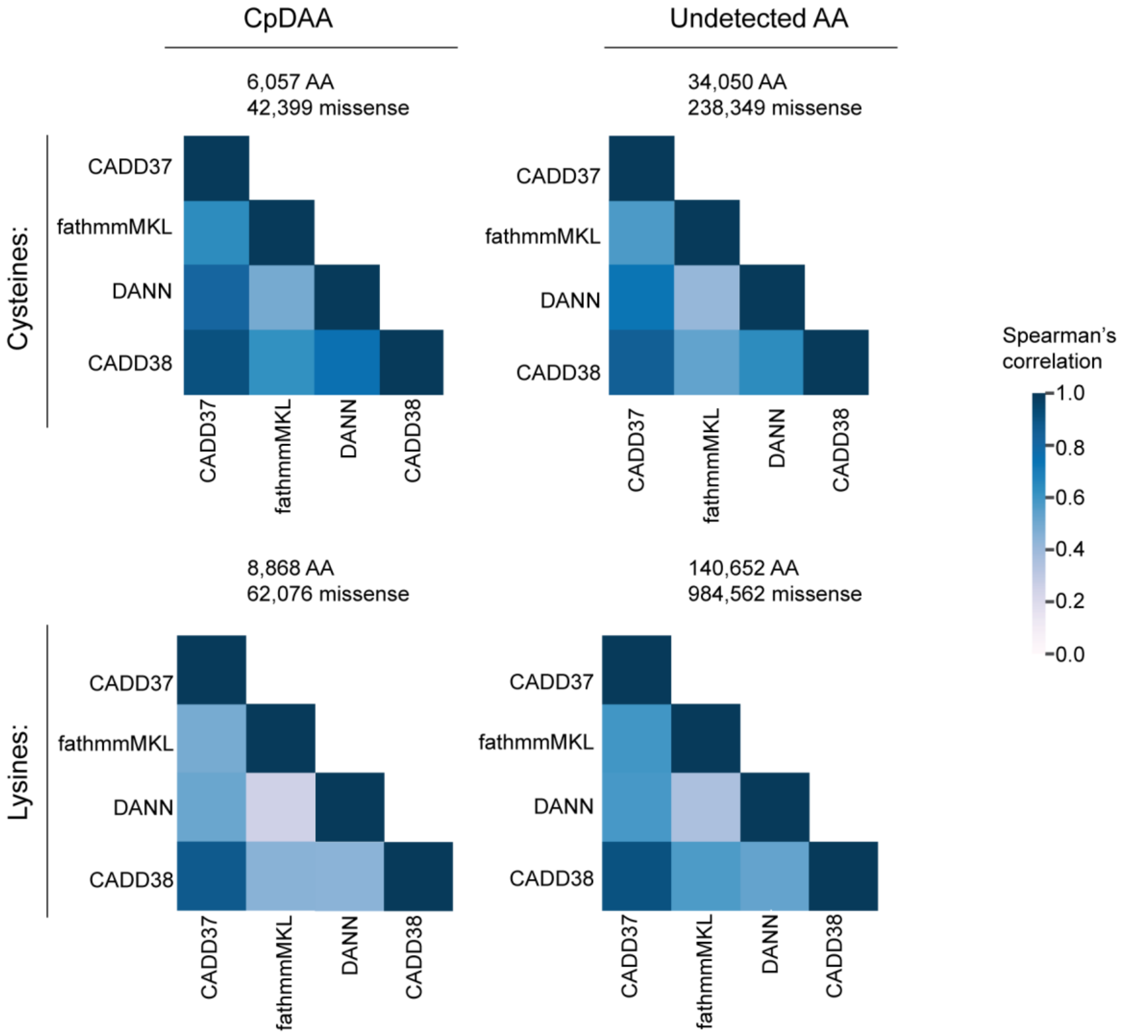


**Figure EV7.** Correlation of machine learning pathogenicity scores for CpD and undetected cysteine and lysine residues. Heatmap color based on Spearman rank correlation coefficient (r). Number of unique amino acid positions and number of possible nonsynonymous variants scored at cysteine and lysine codon coordinates are shown for 3,840 UniProtKB proteins.

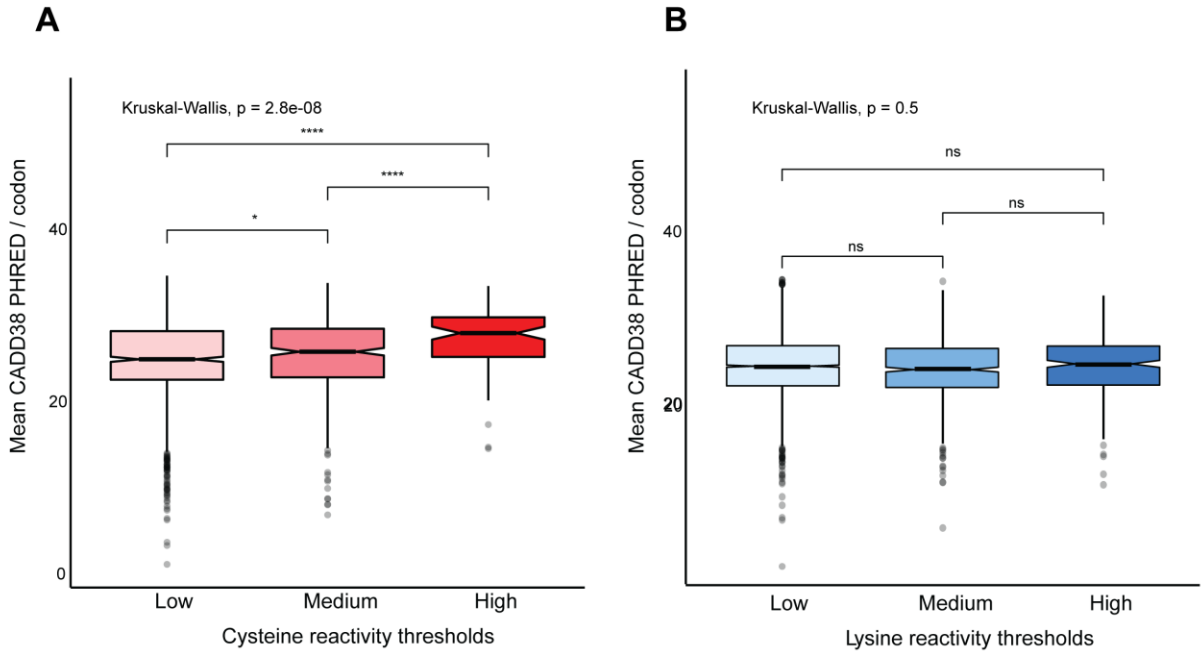


**Figure EV8.** Association between amino acid reactivity and CADD score. A-B) Distribution of the mean CADD38 (model for GRCh38) PHRED score/codon for (A) cysteine (n=1,401) and (B) lysine (n=4,363) CpDAAs of low, medium, and high intrinsic reactivities, defined by isoTOP-ABPP ratios, low ($R_{10:1}$>5), Medium (2<$R_{10:1}$<5), High ($R_{10:1}$<2)(Weerapana et al. 2010; Hacker et al. 2017)}. Kruskal-Wallis test for multiple pairwise-comparisons. Wilcox test used for pairwise comparisons. *, $p$ < 0.1, ****, $p$ < 0.00001.

### REFERENCES

Hacker, S. M., K. M. Backus, M. R. Lazear, S. Forli, B. E. Correia, and B. F. Cravatt. 2017. “Global Profiling of Lysine Reactivity and Ligandability in the Human Proteome.” *Nature Chemistry* 9 (12): 1181–90.

Weerapana, E., C. Wang, G. M. Simon, F. Richter, S. Khare, M. B. Dillon, D. A. Bachovchin, K. Mowen, D. Baker, and B. F. Cravatt. 2010. “Quantitative Reactivity Profiling Predicts Functional Cysteines in Proteomes.” *Nature* 468 (7325): 790–95.
